# How good is Generative Diffusion Model for Enhanced Sampling of Protein Conformations Across Scales and in All-atom Resolution?

**DOI:** 10.1101/2025.01.16.633470

**Authors:** Palash Bera, Jagannath Mondal

## Abstract

Molecular dynamics (MD) simulations are fundamental for probing the structural dynamics of biomolecules, yet their efficiency is limited by the high computational cost of exploring long-timescale events. Generative machine learning (ML) models, particularly the Denoising Diffusion Probabilistic Model (DDPM), offer an emerging strategy to enhance conformational sampling. In this study, we evaluate the capabilities and limitations of DDPM in generating atomistically accurate conformational ensembles across proteins of varying size and structural order, ranging from the 20-residue folded Trp-cage and 58-residue BPTI to the 83-residue intrinsically disordered region Ash1 and the 140-residue intrinsically disordered protein *α*-Synuclein. Training DDPM on relatively short MD trajectories using both torsion angle and all-atom coordinate data, we demonstrate that it can reproduce key structural features such as secondary structure, radius of gyration, and contact maps, while effectively sampling sparsely populated regions of the conformational landscape. Notably, DDPM can also generate novel conformations, including transitions not explicitly observed in the training data. However, the model occasionally overlooks low-probability regions and may produce conformers with unclear physical relevance, warranting independent validation. These limitations are particularly evident in flexible systems such as IDPs. Overall, this work benchmarks DDPM as a viable tool for augmenting MD simulations, offering enhanced sampling with significant computational savings, while noting its limitations in capturing low-populated conformers. At the same time, it highlights the importance of rigorous validation and thoughtful interpretation when deploying generative models in computational biophysics.

## INTRODUCTION

Understanding and predicting the conformational landscapes of proteins are pivotal for unraveling their functions, stability, and interactions within biological systems. While structured proteins adopt well-defined three-dimensional (3D) conformations that can be effectively characterized through experimental and computational methods, intrinsically disordered proteins (IDPs) and intrinsically disordered regions (IDRs) lack such static structures. Instead, IDPs and IDRs exhibit vast conformational ensembles, which are critical for their diverse functional roles, including signaling, regulation, and molecular recognition.^1–4^ Notably, IDPs are implicated in numerous human diseases,^5–9^ such as neurodegenerative disorders and cancers, emphasizing the importance of understanding their long-timescale structural dynamics.

Despite significant advances in experimental and computational approaches, classical molecular dynamics (MD) simulations remain the gold standard for generating conformationally dynamic ensembles of biomolecules at atomic resolution. However, the extensive computational resources required for MD simulations pose a major challenge to the realistic exploration of long-timescale atomistic conformational ensembles. While specialized hardware platforms have enabled sampling of conformational landscapes,^10–12^ their limited accessibility to the broader research community and the high computational demand for studying IDPs and IDRs have restricted their widespread use.

To address these limitations, various enhanced sampling methods have been developed. Techniques such as metadynamics,^13,14^ umbrella sampling,^15,16^ steered MD,^17,18^ and temperature-accelerated MD^19,20^ introduce biases along predefined reaction coordinates to facilitate the sampling of rare events. Alternatively, adaptive sampling^21,22^ utilizes order parameters to guide unbiased simulations, thereby enhancing the exploration of under-sampled regions. Another category of methods, including replica exchange MD (REMD),^23,24^ accelerated MD,^25,26^ and weighted-ensemble simulations^27,28^ enhances sampling by modifying the system’s global Hamiltonian, allowing energy barriers to be surmounted and diverse conformational states to be explored. While these methods have achieved notable success, they often rely on prior knowledge of reaction coordinates or incur significant computational costs.

Recently, machine learning-based approaches have emerged as powerful tools for enhanced sampling. Techniques such as VAMPnets^29^ and time-lagged autoencoders^30^ have been used to extract meaningful collective variables (CVs) from simulation data. Statistical methods, including Markov state models (MSMs)^31–33^ and Hidden Markov Models (HMMs)^34–36^ discretize continuous molecular trajectories into discrete states, providing insights into long-timescale dynamics. However, these approaches often require labor-intensive state-space discretization and struggle with sparse sampling of rare or inaccessible states, necessitating user-guided adaptive sampling.

In addition to traditional and machine learning-guided approaches, generative deep learning models have transformed the study of molecular dynamics and structural prediction. Autoencoder^37–39^ and their variants, such as Variational Autoencoder (VAE),^40,41^ have been utilized to uncover low-dimensional latent representations of complex molecular systems. They have also been used to reconstruct and generate conformation of biomolecules.^42–44^ Generative Adversarial Network (GAN)^45,46^ and Normalizing Flows^47–49^ have shown promise in generating realistic molecular structures and conformations by learning high-dimensional probability distributions. Boltzmann Generators,^50^ which combine generative modeling with statistical mechanics, directly sample equilibrium distributions, enabling efficient exploration of thermodynamic states. Language models, including Long Short-Term Memory (LSTM)^51–54^ and Transformers,^55,56^ have advanced the prediction of state-to-state transitions by capturing temporal dependencies and long-range interactions. Diffusion-based models, such as the Denoising Diffusion Probabilistic Model (DDPM),^57,58^ have emerged as powerful tools for generative tasks, often rivaling or surpassing other generative AI models in producing high-quality images.^59^ DDPM learn the underlying probability distribution of complex data by gradually adding noise to the data (the forward diffusion process) and then learning to reverse this process (the denoising step) to recover the original data distribution. This iterative noising-denoising procedure allows DDPMs to generate high-quality, physically plausible samples, even in high dimensional spaces such as those encountered in biomolecular conformational landscapes.^60–64^

Recent breakthroughs, such as AlphaFold^65,66^ and RoseTTAFold,^67,68^ have revolutionized protein structure prediction. However, while these methods excel at predicting static structures, they cannot generate dynamic conformational ensembles, especially for IDPs, which lack stable structures and exhibit extensive conformational heterogeneity. The potential application of generative artificial intelligence in enhancing sampling has been an active area of discussion in the computational biophysics community.^69^ In particular, the integration of artificial neural network-based approaches with molecular dynamics (MD) simulations has garnered significant attention as a means to improve sampling efficiency and capture rare conformational transitions.^70^ Beyond static structure prediction, the AI-driven generation of Boltzmann-weighted structural ensembles of biomolecules has emerged as an important and rapidly evolving research direction, offering new possibilities for exploring conformational landscapes with improved accuracy and efficiency.^71^ Generative models like GANs and DDPM have recently been applied to explore conformational ensembles of folded proteins and IDPs.^72,73^ Nevertheless, most studies on IDPs rely on coarse-grained simulations, limiting their atomistic detail. Generating all-atom conformational ensembles for larger proteins, particularly IDPs, remains a significant challenge.

In this study, we investigate the potential of DDPM for enhanced sampling and conformational ensemble generation across diverse biomolecular systems. By training DDPM on relatively short MD simulation segments of torsion angles and all-atom coordinates, we evaluate its ability to enhance conformational sampling and accurately reproduce key structural features, including secondary structure, radius of gyration, and contact maps. Our analysis encompasses both folded and intrinsically disordered proteins across a wide range of sequence lengths, specifically the 20-residue Trp-cage mini-protein, the 58-residue folded protein BPTI, the 83-residue intrinsically disordered region (IDR) Ash1, and the 140-residue intrinsically disordered protein (IDP) *α*-Synuclein, which is associated with neurological disorders.

Recent studies have reported improved performance of generative diffusion models for protein conformational sampling by incorporating features such as coarse-grained representations, implicit solvent models, and thermodynamic or sequence-aware conditioning.^72–76^ While these strategies enhance generalization and chemical diversity, our present study deliberately avoids such assumptions. Instead, we focus on evaluating the baseline performance of unconditioned DDPMs trained solely on limited, all-atom MD data across diverse proteins. This minimalistic design allows for a transparent and unbiased benchmark of what diffusion models can achieve without architectural or prior-driven advantages. We believe this distinction is essential for disentangling the effects of model expressivity from those of training data quality or external constraints, and for motivating future extensions toward transferable and physically grounded generative frameworks.

We find that DDPM effectively learns the underlying distributions of the training data, generating thermodynamically consistent conformations that align well with those obtained from significantly longer simulations. Furthermore, DDPM is capable of generating configurations in sparsely populated regions of the free energy landscape, although its ability to fully capture all such low-probability states remains limited. In certain cases, the model omits rare conformers, while occasionally generating novel states whose physical significance requires independent validation.

## METHODS

### Theoretical framework of the Denoising Diffusion Probabilistic Model (DDPM)

The DDPM is an ML-based generative model that is formulated by inspiring the principles of real diffusion processes.^57,58^ After training, the DDPM can generate new samples starting from purely random noise. These generated samples closely follow the statistical probability distribution of the training data, making them physically realistic and representative of the original dataset.

### Forward and Reverse Processes in DDPM

The DDPM operates through two complementary processes: (i) the forward or noising process, and (ii) the reverse or denoising process. In the forward process, noise is progressively added to the data over multiple timesteps, eventually converting it into pure Gaussian noise. Mathematically, if *Q*(*x*_0_) represents the real data distribution, the forward-nosing process can be described as a Markovian diffusion process with the following transition probabilities:

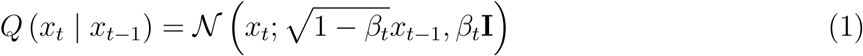

Here, *t* is the timestep, *t* ∈ [0*, T* ], T denotes the total number of timesteps, *x_t_* represents the noisy data at timestep *t*, and *β_t_* is the predefined noise schedule which controls the amount of noise added between timesteps *t* − 1 and *t*. Using the reparameterization trick, we can rewrite equation 1 in a closed form as:

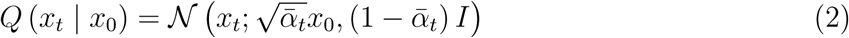

where *α_t_* = 1 − *β_t_*, and 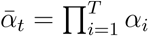. This reparameterization trick helps us to sample at any arbitrary timestep t (see subsection *Brief overview of reparameterization scheme in DDPM* in the supplementary information (SI) for details).

On the other hand, the reverse process uses a neural network trained to model the reverse of the noising process by predicting the noise added at each step, effectively reconstructing the original data from the noisy input. The reverse denoising process is mathematically defined as:

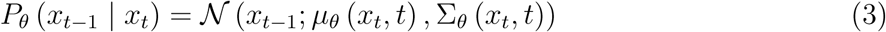

where *µ_θ_* and Σ*_θ_* represent the learnable mean and variance of the conditional distribution, parameterized by the neural network.

### Model Architecture and Hyperparameters

In this work, we employ a commonly used architecture for this task, namely U-Net, an encoder-decoder neural network originally introduced by Ronneberger et al.^77^ U-Net also has been widely used for tasks such as medical image segmentation.

Figure 1 represents a schematic overview of the noising and denoising processes. In our study, the input data at timestep *t* = 0 consist of either the backbone torsion angles (*φ* and *ψ*) or the coordinates of all heavy atoms in the proteins (excluding hydrogen atoms). We set the total timesteps for the diffusion process to *T* = 1000. The lower panel of the figure 1 illustrates the U-Net model used in our study, particularly when the input data are backbone torsion angles. The U-Net architecture primarily involves downsampling and upsampling operations. Each downsampling stage comprises ResNet blocks combined with a downsampling operation (max-pooling). Similarly, each upsampling stage consists of ResNet blocks with upsampling operations (transposed convolutions). When using heavy-atom coordinates as input data, we observed that the model required a greater number of downsampling and upsampling blocks compared to backbone torsion angles to generate realistic protein conformations. This is likely due to the significantly larger feature space represented by the coordinates, requiring additional layers for the diffusion model to accurately learn the underlying distribution of the training data. A comprehensive description of the U-Net architectures used for the various systems in our study is provided in the SI (see subsection *Adapting U-Net Architectures Based on Input Feature Complexity* in the SI). To optimize the neural network parameters (weights and biases), we employed the Adam optimizer^78^ with a learning rate 1 × 10*^−^*^5^, minimizing the mean squared error (*L*_2_ loss) between the predicted and added noise. Additionally, an exponential moving average (EMA) with a decay rate of 0.995 was applied to enhance the stability of the optimizing process. The code and comprehensive documentation for training the DDPM can be accessed on GitHub via the following link: https://github.com/palash892/DDPM_conformational_ensemble

**Figure 1:**
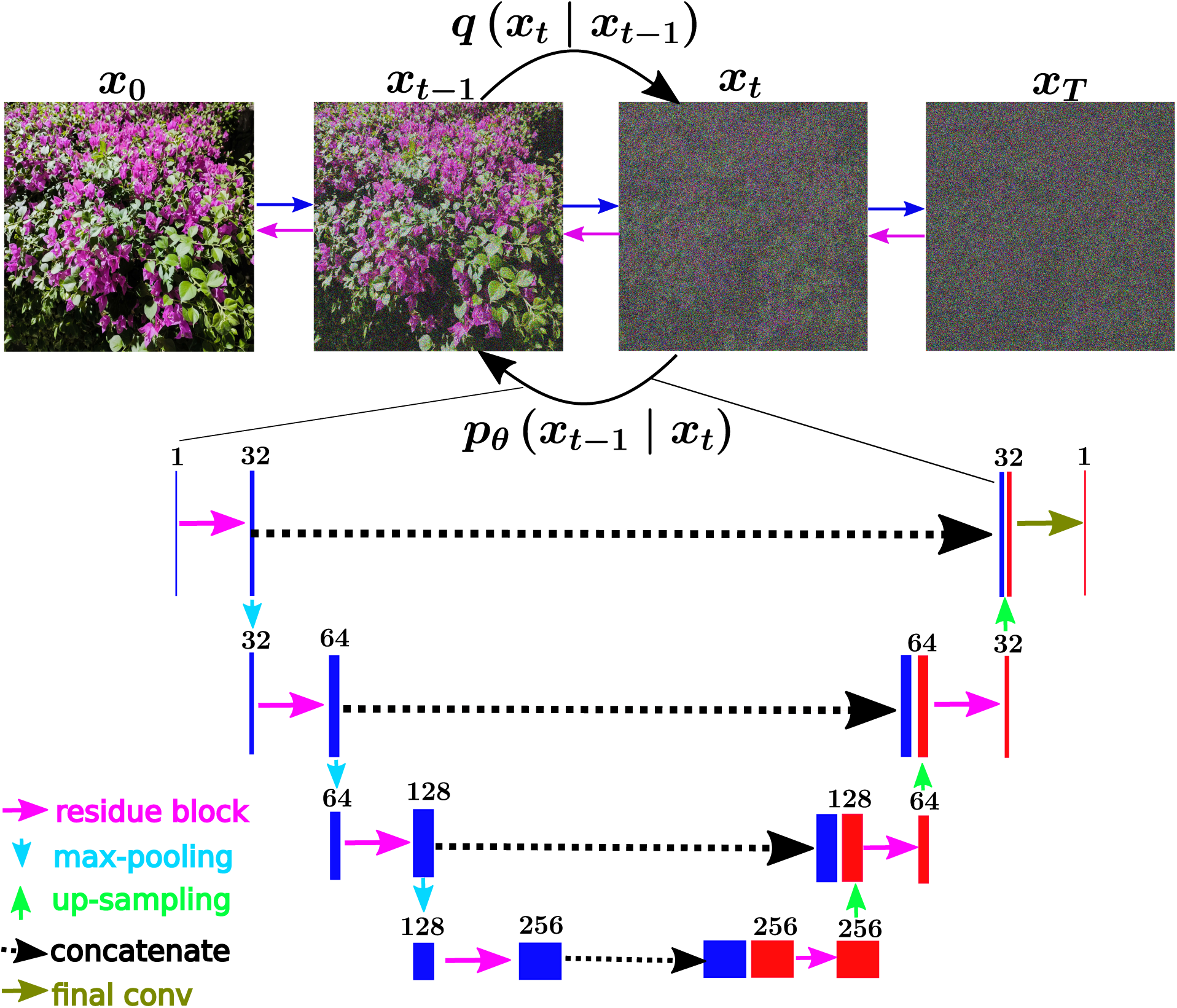
A schematic overview of the noising and denoising processes. We illustrated the noising-denoising processes using an image of a Paperflower clicked at the TIFRH campus during summer. The upper panel depicts the forward (noising) process, represented by the blue arrow, where the original input image *x*_0_ is progressively corrupted by adding random noise, and the reverse (denoising) process, represented by the black arrow, where a neural network predicts the added noise. The lower panel illustrates the U-Net model architecture. The U-Net consists of downsampling and upsampling operations, where each downsampling stage features residue or ResNet blocks with max-pooling, and similarly each upsampling stage includes residue or ResNet blocks with transposed convolutions.

### Training datasets

For a fair and rigorous assessment of the generative diffusion models in enhancing conformational sampling, we made a call to go beyond putative toy-models and model biomolecules such as Alanine dipeptides and its variants. Nonetheless, these have been extensively explored by previous effort.^74^ In our study, we employed MD simulation datasets provided by D. E. Shaw Research^10,79^ for four distinct biomolecular systems, encompassing both folded proteins and IDPs. Our choice ensured that both fold complexity (mini protein to disordered protein) and variation in sequence lengths (ranging between 20 to 140) are taken into account during our investigations. These systems are detailed below:

- **Trp-cage Mini-Protein:** For the folded mini-protein Trp-cage, we employed 100 *µs*-long trajectories as ‘ground truth’ data. This globular 20-residue protein is frequently used as a benchmark system for computational studies.
- *α***-Synuclein:** *α*-Synuclein, a 140-residue IDP, is characterized by its significant conformational heterogeneity. The MD simulation data spanning 73 *µs* were used.
- **BPTI (Bovine Pancreatic Trypsin Inhibitor):** This 58-residue folded protein was selected to test the DDPM’s ability to generate all-atom conformational ensembles. We used 10 *µs* of unbiased MD simulation data.
- **Ash1:** The IDR Ash1 consists of 83 residues. Its conformational heterogeneity was studied using 30 *µs* of unbiased MD simulation data.

Each of the parent simulation data for each system was obtained from one single continuous extensively long MD simulation, rather than ensemble aggregated multi-trajectory simulations. For all of the systems, we decided to consider the leading segment of the simulation trajectory as training data. The percentage of data used for training the model varies across different systems and was determined based on the minima of their respective free energy surfaces along specific CVs. Since DDPM is a probabilistic model that learns the probability distribution of the training data, this approach ensures that the training data encompasses major relevant minima. However, these training data points are sparse compared to the full dataset. For example, in the case of Trp-cage, the first 10% (50,000 frames) of the total simulation data was sufficient for training, effectively capturing the essential features of the system. However, for the more complex *α*-Synuclein, first 50% (36,562 frames) of the simulation data was necessary to achieve similar results, in case. Notably, using only 10% of the *α*-Synuclein trajectory as training data did not work well, in the sense that this portion of the actual MD trajectory failed to cover all the major basins in the free energy surface (FES). We further analyzed other portions of the trajectory, specifically 20%, 30%, and 40%, and compared their FES plots (see Figure S1). These too were insufficient in capturing the full conformational landscape. However, the first 50% of the trajectory included all key minima of the FES, albeit sparsely populated, and was thus chosen as the training set. This limitation was attributed to insufficient frame saving in the MD simulation and the intrinsic complexity of the system, which demands a more comprehensive dataset for the model to perform effectively. In particular the saving frequency of the original DESRES simulation trajectory was significantly lower in case of *α*-Synuclein than that in case of Trp-cage.

## RESULTS

Our central objective in this study was to systematically evaluate the potential of denoising diffusion probabilistic models (DDPMs) as a tool for enhanced sampling of biomolecular conformational landscapes. To this end, we selected a diverse set of protein systems spanning a broad spectrum of structural complexity, including from compact, well-folded globular proteins such as Trp-cage and BPTI to highly dynamic intrinsically disordered proteins (IDPs) like *α*-Synuclein and Ash1. In order to rigorously assess the generative capabilities of DDPMs without the influence of system-agnostic priors or inductive biases, we employed system-specific training for each protein. This design choice enabled us to directly quantify how well DDPMs can reconstruct conformational ensembles and recover key thermodynamic features from limited molecular dynamics (MD) data. The following subsections present a detailed case-by-case analysis of this evaluation.

### Sampling Accuracy of DDPM on Benchmark Protein Systems along dihedral space

To understand the sampling accuracy of the DDPM, we first focus on two benchmark systems: the globular 20-residue mini-protein Trp-cage and the 140-residue IDP *α*-Synuclein. As input data for training the DDPM, we selected the backbone dihedral angles (*φ, ψ*) of these proteins. This resulted in a total of 38 torsion angles for Trp-cage and 278 for *α*-Synuclein. Consequently, the input data vectors are represented as: 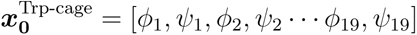 and 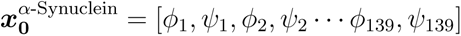 corresponding to the Trp-cage mini-protein and *α*-Synuclein, respectively. To ensure a fair assessment of DDPMs true capability in augmenting sampling, we deliberately avoided generating training data by randomly reshuffling segments of simulation trajectories, as this would introduce bias by a priori incorporating knowledge of the full trajectory. Instead, we consciously chose to use only the leading segment of the simulation trajectories as training data for DDPM. This approach better reflects a realistic scenario in which a simulator, starting from an initial configuration, has sampled only a limited trajectory length and subsequently relies on generative machine learning to extend sampling for improved conformational space exploration.

### 2D Free Energy Surfaces

To visualize the high dimensional torsion angles of these systems, we performed a principal component analysis (PCA) on the torsion angles and plotted the free energy surface (FES) using the first two principal components (PC-1 and PC-2). Figures 2(a) and (d) present the 2D FES plots along PC-1 and PC-2 for Trp-cage and *α*-Synuclein, respectively. To examine the capability of DDPM for enhanced sampling, we trained the model on subsets of the simulation data instead of the entire time series. For Trp-cage, we used the first 10% (50000 frames) of the total simulation data, while for *α*-Synuclein, we selected the first 50% (36562 frames) . Using these subsets, we generated 2D FES plots along PC-1 and PC-2 (Figures 2(b) and (e)). These plots indicate the sparsity in the free energy surface compared to those derived from the full 100% datasets. We then trained the DDPM on these selected subsets and generated the same number of samples as in the original 100% simulation datasets. The generated dihedral angles were projected onto the principal components defined by the original MD data. Figures 2(c) and (f) depict the corresponding FES plots for Trp-cage and *α*-Synuclein. Quite interestingly, the DDPM-generated FES plots closely resemble those derived from the full 100% simulation datasets. More importantly, the sparse regions observed in Figures 2(b) and (e) were effectively sampled by the data generated by DDPM. But a closer comparison of DDPM-generated landscape with that of 100% MD-data suggests that ML-generated landscape also misses a minor yet non-negligible population, particularly the shoulder around (PC1, PC2) = (-4,-2). For Alpha-synuclein as well, while the DDPM-generated conformations resolves the FES to a reasonable extent, the overall DDPM-generated FES is significantly more smoother than its MD simulation counterparts. One possible interpretation might be that DDPM might either miss out sampling the low-probability regions not present in training data (as in case of Trp-cage) or smooth out the FES via its inherent interpolation of transition between sparse regions (as in case of *α*-Synuclein). Collectively, these results suggest that DDPM reasonably learns the distribution of the training data, enabling it to generate thermodynamically consistent torsion data, while certain low-probability regions not present in training data not being replicated by it .

**Figure 2:**
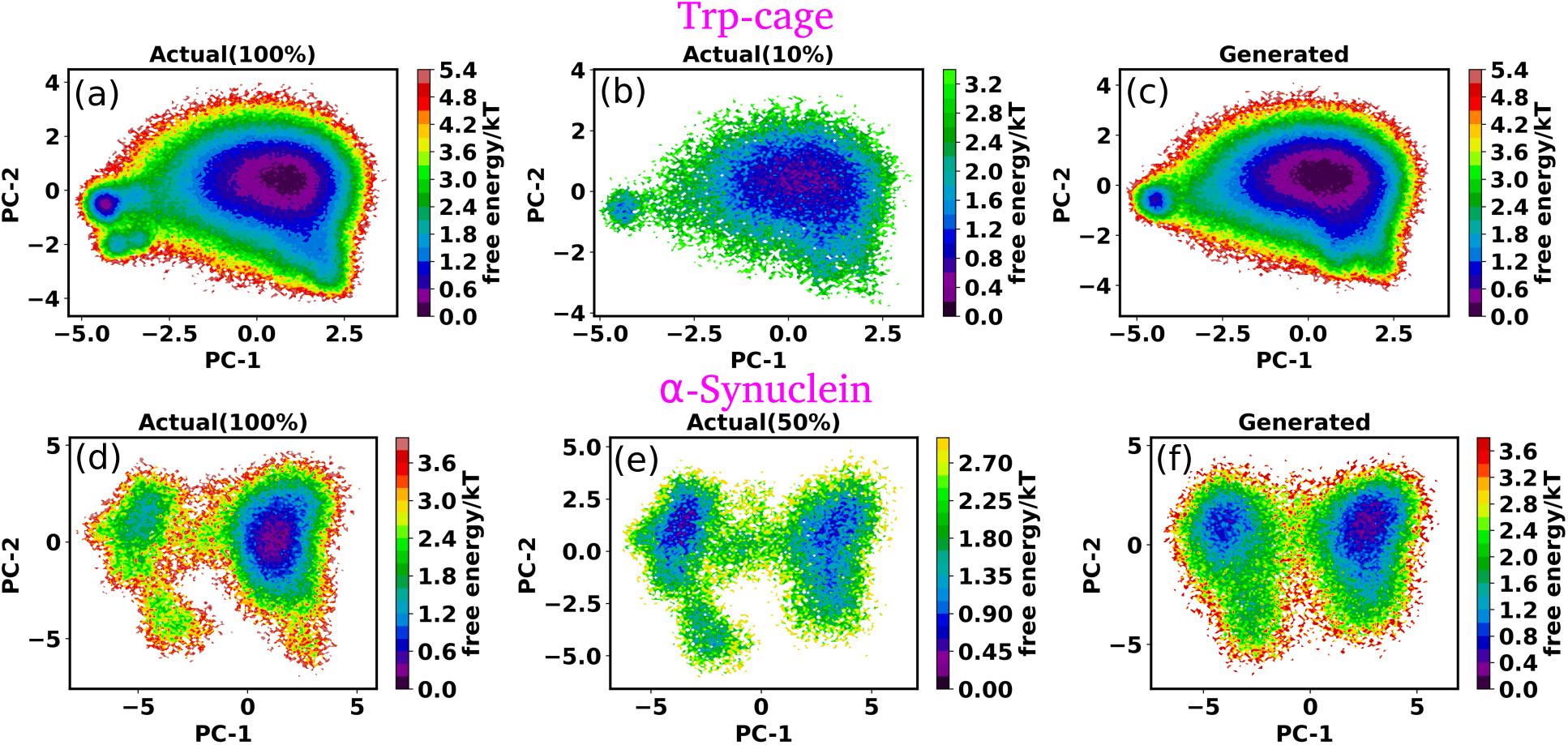
The free energy surfaces (FES) of Trp-cage and *α*-Synuclein based on principal component analysis (PCA) of torsion angles. (a) and (d) show the 2D FES plots along the first two principal components (PC-1 and PC-2) derived from the full molecular dynamics (MD) simulation datasets for Trp-cage and *α*-Synuclein, respectively. (b) and (e) depict the FES plots using subsets of the simulation data, corresponding to 10% of the total data for Trp-cage and 50% for *α*-Synuclein. These subsets highlight the sparsity of the sampled free energy surfaces compared to the full datasets. (c) and (f) present the FES plots generated from the DDPM-trained models, which were trained on the reduced subsets. Remarkably, the DDPM-generated FES plots closely resemble those from the full datasets and effectively sample the sparse regions observed in (b) and (e).

#### Characterizing the DDPM-augmented Enhanced Sampling

For a close investigation of the enhanced sampling, we analyzed the Ramachandran plots of specific *φ* − *ψ* combinations for both systems. We plotted both the full simulated data, selected subsets as previously described, and DDPM-generated data. Figures 3(a) and (b) show the representative Ramachandran plots of (*φ*_6_ − *ψ*_6_) and (*φ*_17_ − *ψ*_17_) for Trp-cage mini-protein, while Figure 3(c) depicts the plot of (*φ*_5_ − *ψ*_5_) for IDP *α*-Synuclein. Additional combinations of *φ* − *ψ* for both systems are provided in the Supporting Information (SI) (Figures S2 and S3). For *α*-Synuclein, which has 139 possible consecutive *φ* − *ψ* combinations, the first 5 combinations are presented in the SI. Our analysis reveals two key features for enhanced sampling: (i) The DDPM can effectively enhance sampling in regions where training data is sparsely distributed, as highlighted by the red boxes in Figures 3(a) and (c). These regions, which are sparsely populated in the training data, are significantly better sampled in the DDPM-generated data. (ii) The DDPM can generate new samples with very low probabilities, absent in the training data but present in the full 100% simulation dataset. This behavior is illustrated in Figure 3(b), where the highlighted region in the DDPM-generated data corresponds to previously unsampled areas of the Ramachandran plot. These observations are also consistent with the previous study by Wang et al.^74^ Nevertheless, our findings indicate DDPM’s ability to enhance sampling through both interpolations of existing data distributions and generation of previously unobserved samples. On similar note, we also found cases where DDPM initiates transitions connecting two spatially well-separated basins, otherwise unobserved in even 100 % datasets (Figure 3b)). However, independent validations would be required for assessing the significance of these newly generated low probability regimes which DDPM might be ‘dreaming of’, as has been previously noted. ^74^

**Figure 3:**
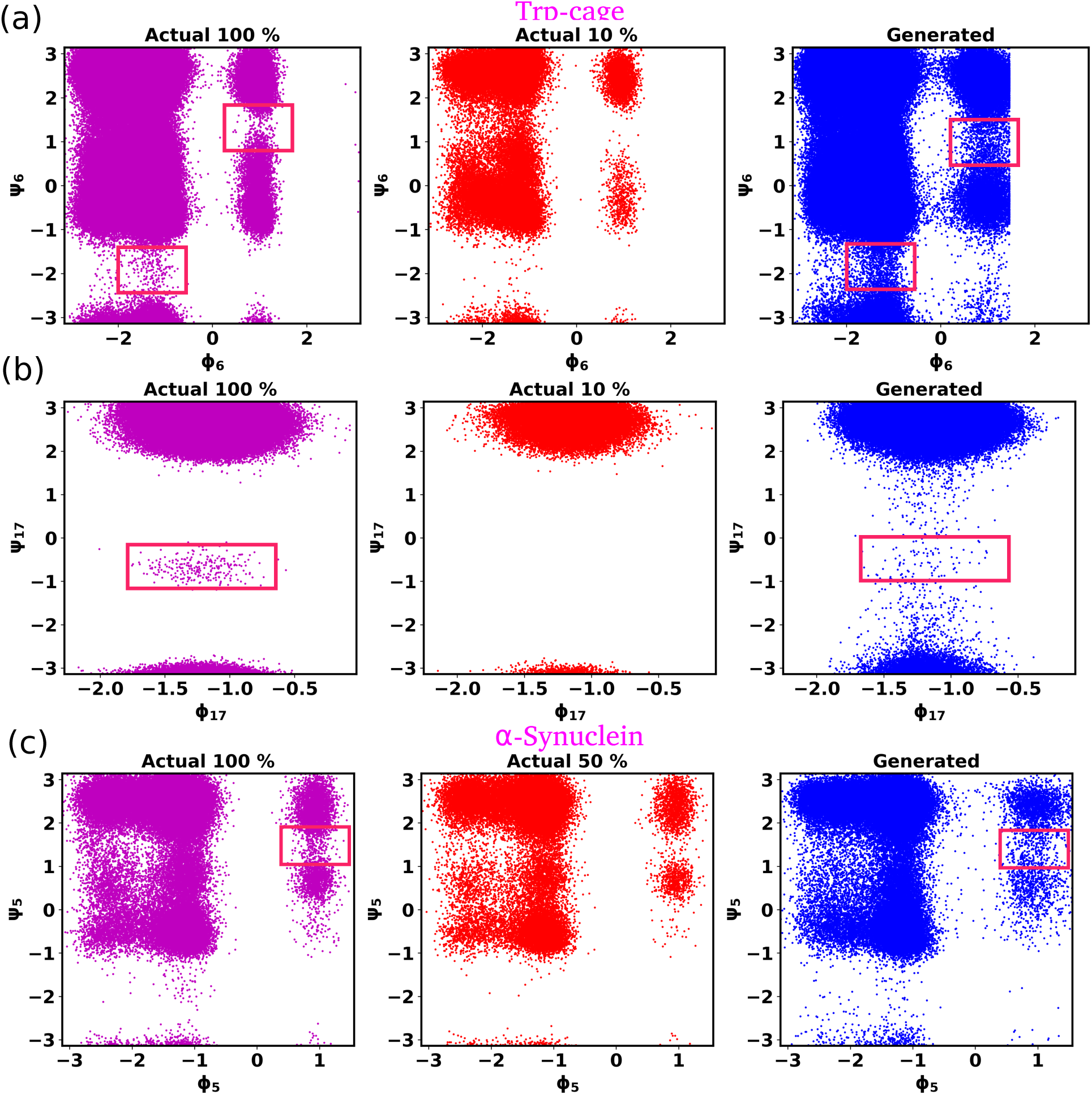
Ramachandran plots showing how DDPM improves sampling coverage. (a) and (b) show the Ramachandran plots for the *φ*–*ψ* combinations (*φ*_6_ –*ψ*_6_) and (*φ*_17_ –*ψ*_17_) in Trp-cage, respectively. (c) presents the plot for the *φ–ψ* combination (*φ*_5_–*ψ*_5_) in IDP *α*-Synuclein. We plotted the complete simulated data, the selected subsets for training (10% and 50%), and the DDPM-generated data. The red boxes in (a) and (c) (Generated) highlight regions with sparse sampling in the training data that are significantly enhanced in the DDPM-generated data, illustrating interpolation capabilities. In (b) (Generated), the red box highlights new low-probability samples generated by DDPM, absent in the training data but present in the full 100% simulation dataset.

#### Probability Distributions of Torsion Angles

To further validate DDPM’s performance, we compared the individual probability distributions of torsion angles *φ* and *ψ*. For this comparison, we chose the complete simulated data, a subset of simulated data, and DDPM-generated data, as previously outlined. Figures 4(a-d) depict the probability distributions of selected dihedral angles for Trp-cage mini-protein, while Figures 4(e-f) illustrate these distributions for *α*-Synuclein. Additional distributions for other *φ* and *ψ* angles are reported in the SI (Figures S4-S7). The results indicate that for both systems, the distribution of generated torsion angles closely aligns with that of the original data. Notably, the DDPM captures the bimodal distribution of torsion angles quite accurately, as demonstrated in Figures 4(b) and (f). This accuracy highlights DDPM’s ability to learn and generate complex distributions within molecular dynamics simulations, barring some low probability regimes.

**Figure 4:**
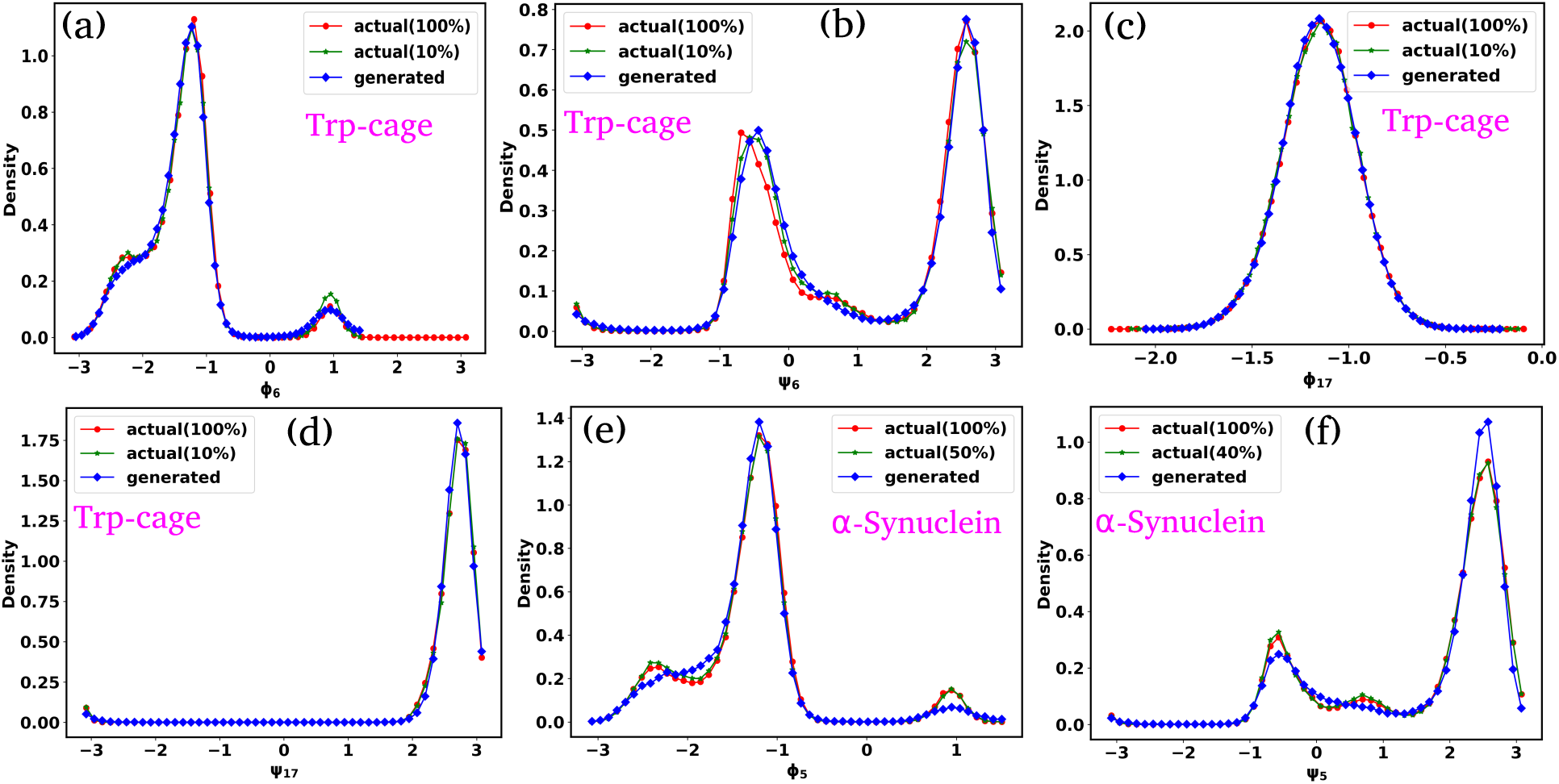
Comparison of individual probability distributions for selected torsion angles *φ* and *ψ*. (a-d) Probability distributions for selected dihedral angles in Trp-cage mini-protein. (e-f) Probability distributions for selected dihedral angles in *α*-Synuclein. The DDPM-generated distributions closely match the original simulated data. More importantly, the DDPM accurately captures the bimodal distribution of torsion angles, ((b) and (f)).

To assess how well the model learns to generate physically meaningful conformations, we quantified the agreement between the model-generated and true torsion angle distributions using the Kullback-Leibler (KL) divergence. The KL divergence is defined as:

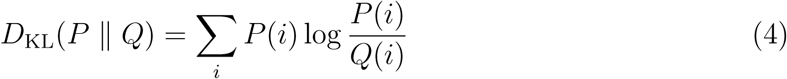

where *P* (*i*) is the true probability of observing torsion angle *i* in the benchmark dataset, and *Q*(*i*) is the probability from the model-generated data. Physically, *D*_KL_ measures how much information is lost when *Q* is used to approximate *P* ; a lower value indicates better agreement between the two distributions.

To evaluate how this divergence evolves with training, we saved the model at fixed epoch intervals and, for each saved model, generated a number of samples equal to the size of the full dataset (100% data). We then computed the KL divergence for each individual torsion angle by comparing the distributions from the true and generated data. The final KL divergence was obtained by averaging these values across all torsion angles.

Figures S8(a) and (b) show the KL divergence as a function of training epochs for Trp-cage mini-protein and *α*-Synuclein, respectively. The plots show a clear monotonic decrease in KL divergence, approaching zero as training progresses. This indicates that the model becomes increasingly effective at reproducing the underlying distribution of torsion angles, providing strong evidence that the generated samples converge toward the true thermodynamic distribution.

### Training DDPM on All-Atom Protein Coordinates

In the preceding sections, we discussed the ability of the DDPM to explore sparsely sampled and previously unsampled regions of the free energy landscape, highlighting its potential as a tool for enhancing sampling in molecular dynamics simulations. However, our previous analyses used backbone torsion angles as input data for training the DDPM, which restricted its output to torsion angles alone. While torsion angles are insightful, they can not fully describe the structural properties of complex biomolecules. To address this limitation, we expanded our approach by training the DDPM using the cartesian coordinates of all heavy atoms as input data. This approach enables the generation of complete conformational ensembles of biomolecules, from which various structural properties can be calculated. In addition to the Trp-cage mini protein and the IDP *α*-Synuclein, we included two additional systems: the 58-residue folded protein BPTI and the 83-residue IDR transcription factor Ash1. Specifically, we selected 40% (19,960 frames) of the BPTI trajectory and 25% (42,500 frames) of the Ash1 trajectory to train the DDPM. For Trp-cage and *α*-Synuclein, we used the same fraction of the initial MD simulation data as in previous analyses. All the aforementioned percentages of the trajectories are selected from the initial portion of the MD simulation data. For DDPM training, we first excluded hydrogen atoms from the trajectories, then centered them at the origin and aligned them with the first frame of each respective trajectory.^44,80^ These transformations ensure that the resulting coordinates are invariant to both rotation and translation. Subsequently, these aligned coordinates were scaled between -1 and +1 to serve as input data for the DDPM. After generating coordinates with the DDPM, these outputs were scaled back to their real-space values across all frames.

### Evaluation of Conformational Ensembles for Folded Proteins

#### Validation Using Trp-cage Mini Protein

To evaluate the structural accuracy of the conformational ensembles generated by the DDPM, we focused initially on two folded proteins: Trp-cage and BPTI. We calculated the backbone torsion angles from the DDPM-generated all-atom conformations and then performed PCA on these dihedral angles. Figures S9 show FES plots of the torsion angles along the two principal axes, PC-1 and PC-2. These plots reveal that the FES derived from the DDPM-generated data aligns closely with the original 100% MD simulation data. Furthermore, the generated conformations enhanced sampling in regions sparsely sampled in the training data (10%). These observations are consistent with our previous analyses of FES plots obtained when training the DDPM directly on torsion angle data, reaffirming the model’s ability to enhance sampling effectively.

Next, we analyzed additional structural properties, such as the average pairwise distance and contact probability between residues (*Cα* − *Cα*). Pairwise distances were calculated for *Cα* atoms across all frames, and contact probabilities were determined by defining a threshold of 0.8 nm. A contact between two *Cα* atoms was assigned a value of 1 if their distance was below the threshold, and 0 otherwise. These calculations were performed for each frame and then averaged across all frames. Panels (a) and (b) in Figure 5 represent the distance map and contact map for the Trp-cage mini protein, derived from the original 100% MD simulation data, the subset of training data, and the DDPM-generated data. These maps demonstrate that the DDPM-generated conformations accurately capture the structural features of the protein, showing excellent agreement with the original data while improving sampling beyond the limited training set.

**Figure 5:**
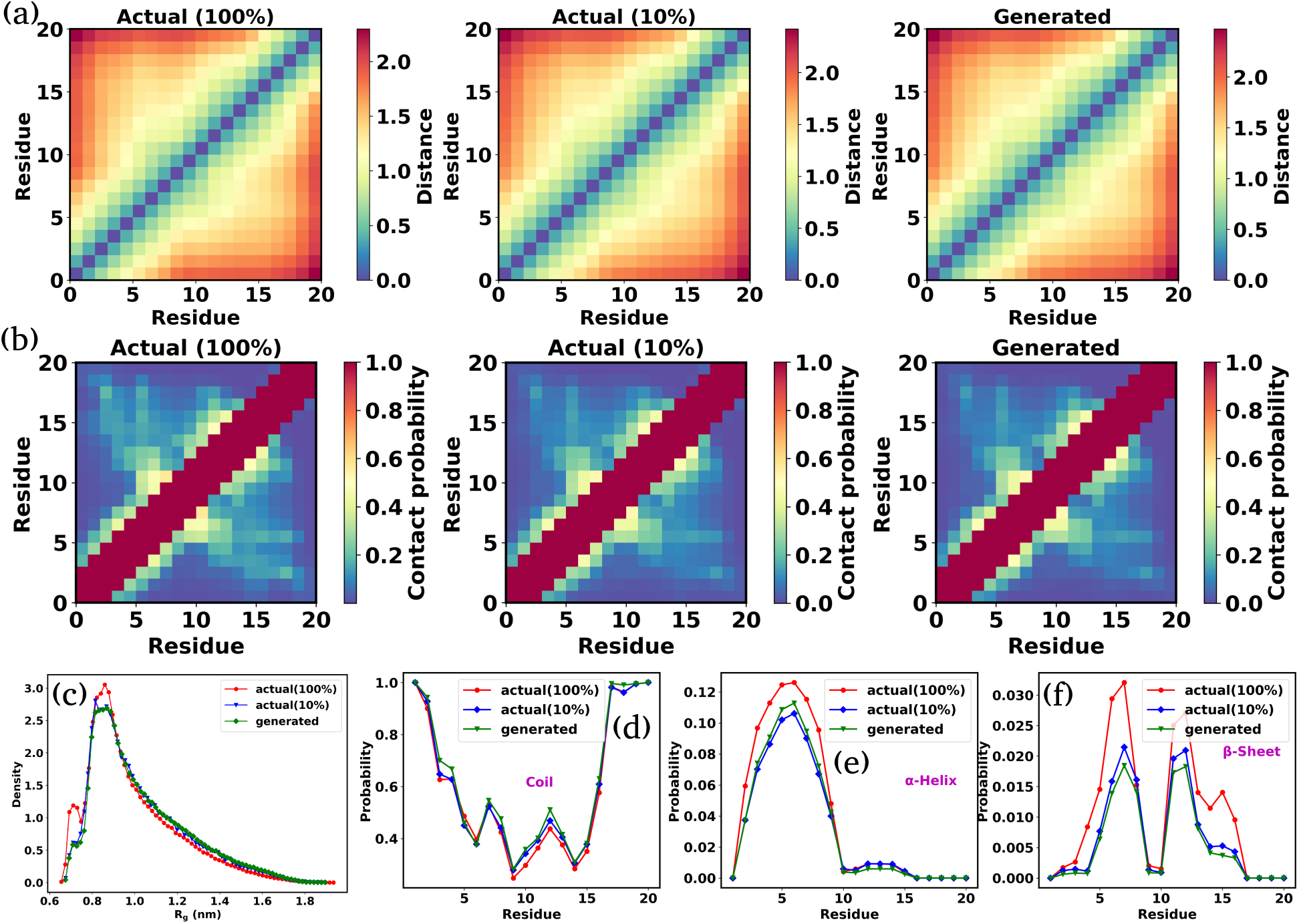
Structural analysis of DDPM-generated conformations for the Trp-cage mini-protein. (a) Distance map and (b) contact map showing pairwise distances and contact probabilities between *Cα* atoms, calculated from the original 100% MD simulation data, the training subset, and DDPM-generated data. (c) Distribution of radius of gyration (*R_g_*) values for the complete 100% MD simulation data, training subset, and DDPM-generated data. (d-f) Secondary structure probability distribution as a function of residue index for Coil, *α*-Helix, and *β*-Sheet, respectively, for the original, training subset, and DDPM-generated conformations. The results suggest that the DDPM-generated ensembles accurately replicate the structural features of the original data, preserving both global compactness and local secondary structure characteristics.

To evaluate compactness and local structural features, we compared the distribution of radius of gyration (*R_g_*) and the secondary structure between the actual and DDPM-generated conformations. We calculated the counts of secondary structures, mainly Coil, *α*-Helix, and *β*-Sheet, for each residue in every frame and converted these counts into probabilities by dividing them by the total number of frames. Figure 5(c) represents the distribution of *R_g_* values for the complete 100% MD simulation data, the training subset, and the DDPM-generated data. Similarly, Figures 5(d-f) show the secondary structure probabilities as functions of residue indices for Coil, *α*-Helix, and *β*-Sheet, respectively. The results indicate that the DDPM-generated ensembles closely replicate the conformational features of the original simulation data, maintaining both global compactness and local secondary structure characteristics with high accuracy. However, certain deviations are observed between DDPM-generated data and fully MD-simulated data, particularly in reproducing the probabilities of *α*-helix and *β*-sheet formations.

#### Validation Using Folded Protein BPTI

To investigate how the performance of DDPM scales across protein sizes (still within folded protein), we also generated the conformational ensemble for the larger 58-residue folded protein BPTI. Backbone torsion angles were calculated from the generated as well as original conformations, followed by PCA to extract principal components. Figure 6(a) presents the FES plots along the first two principal components (PC-1 and PC-2) for the complete 100% MD simulation data, the training subset, and the DDPM-generated data. The generated conformations reproduce trends in the FES that align closely with the original full dataset, while also sampling sparsely populated regions of the training data effectively. Additional comparisons of the distance map, contact map, *R_g_* distribution, and secondary structure (Figures 6(b), (c), (d), and (e-g), respectively) further validate that the DDPM-generated conformations closely match the structural properties of the original data. Notably, the secondary structure plots indicate a probability of 1 for certain residues, confirming that BPTI maintains a rigid and consistent secondary structure. In this regard, it is noteworthy that the residue-wise helical and *β*-sheet propensities in BPTI exhibit a much closer agreement between DDPM-generated conformations and fully MD-simulated data compared to those of Trp-cage, despite the latters significantly shorter sequence length (20 residues). We hypothesize that this discrepancy arises from the inherent flexibility of the Trp-cage fold.

**Figure 6:**
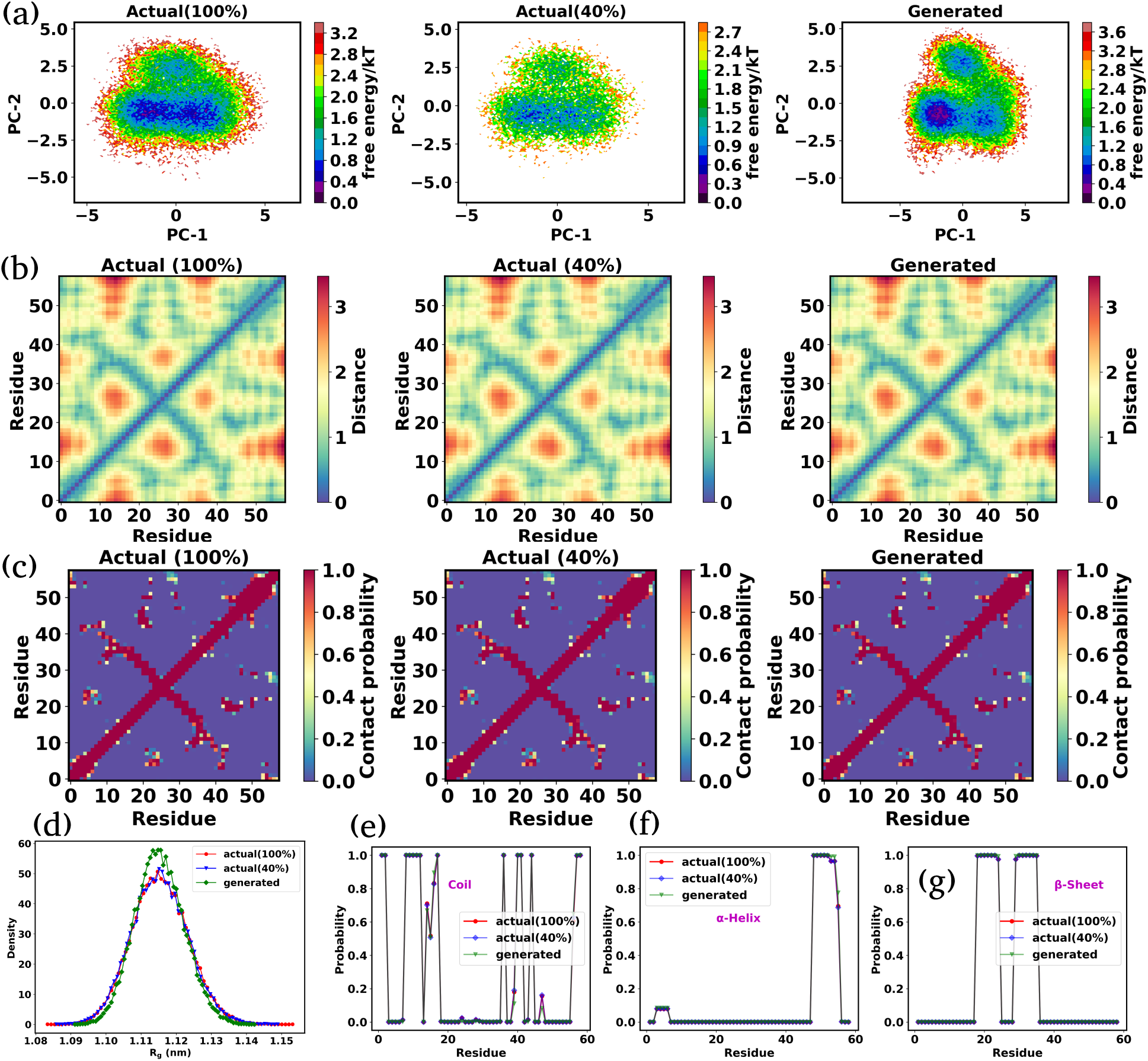
Structural analysis of the DDPM-generated conformational ensemble of folded protein BPTI . (a) FES plots along the first two principal components (PC-1 and PC-2) for the full 100% MD simulation data, the training subset, and the DDPM-generated data, demonstrating that the DDPM-generated all-atom conformations accurately replicate the trends in the FES while effectively sampling sparsely populated regions. (b) Distance map, (c) contact map, and (d) (*R_g_*) distributions, highlighting the structural similarities between the DDPM-generated and original data. (e-g) Secondary structure probabilities for Coil, *α*-Helix, and *β*-Sheet respectively. The secondary structure traces for the full MD and the generated ensembles actually overlap almost perfectly, rendering red trace corresponding to full MD data nearly invisible.

### Evaluation of Conformational Ensembles for IDP and IDR

#### Validation Using IDR Ash1

In the earlier section, we highlighted the capability of DDPM in accurately generating conformations of folded proteins, specifically the 20-residue globular protein Trp-cage and the 58-residue folded protein BPTI. Now, we shift our focus to intrinsically disordered systems, IDRs and IDPs, which lack stable configurations and instead form dynamic and heterogeneous ensembles of conformations. We first examined the 83-residue IDR Ash1 using all-atom molecular dynamics simulations. In a similar fashion to folded protein, backbone torsion angles were calculated from the conformations, and PCA was performed on the torsion angle data. Figure 7(a) shows the 2D FES plots along the first two principal components (PC-1 and PC-2) for the complete MD simulation data, the training subset, and the DDPM-generated data. These plots suggest that the FES plot by the DDPM-generated conformational ensemble of the IDR is qualitatively captured the FES plot of the original data and the sparsely sampled region in the training data is well sampled by the DDPM-generated data. However, certain deviations from the fully MD-simulated dataset also become apparent. Notably, the DDPM-generated landscape exhibits a higher sampling probability in the basin at (PC1, PC2) ≈ (-2,0) compared to the original dataset. with relatively higher probability than the original datasets. This discrepancy appears to be influenced by the quality of training data.

**Figure 7:**
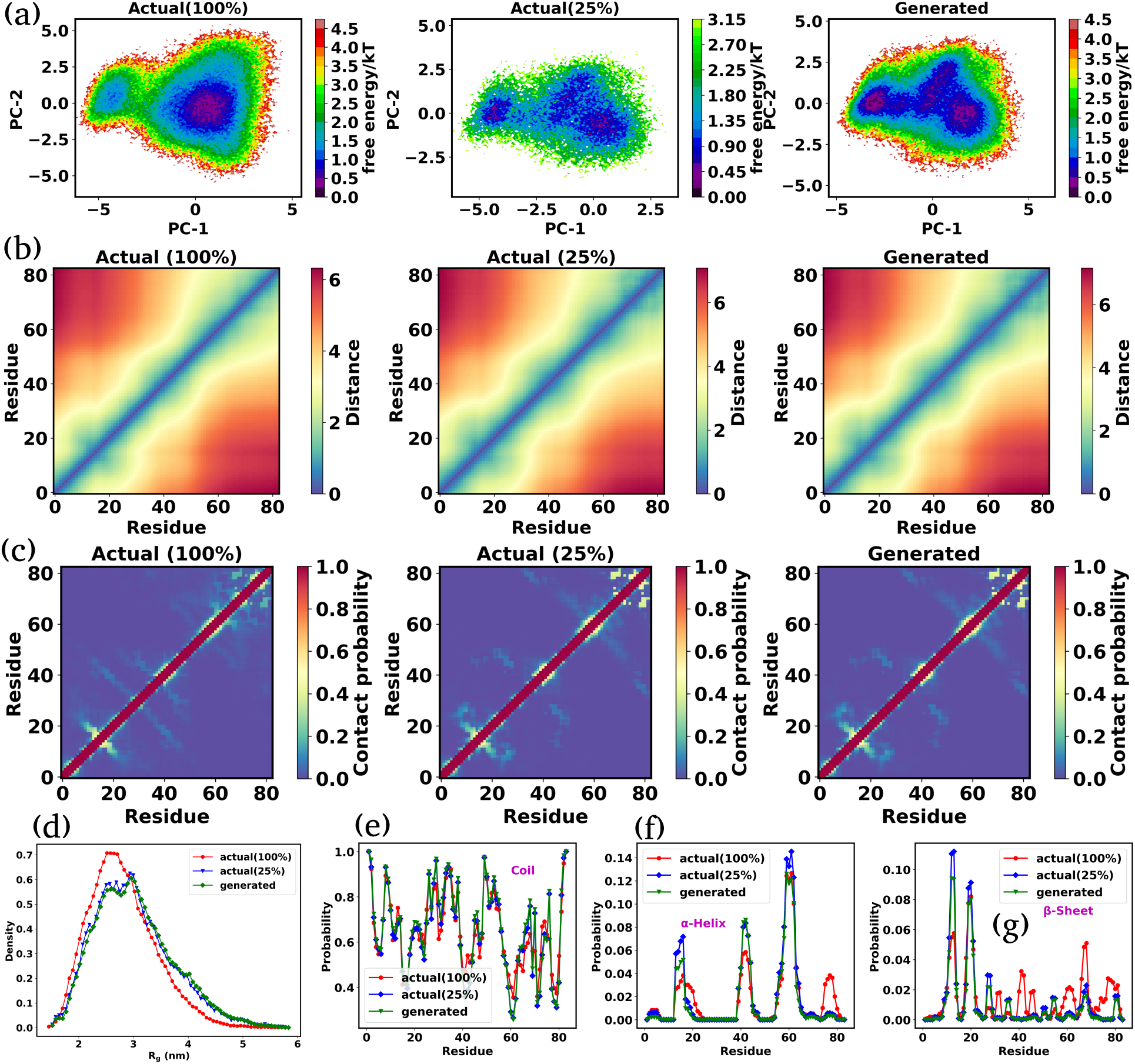
Structural analysis of the intrinsically disordered region (IDR) Ash1. (a) 2D free energy surface (FES) plots along the first two principal components (PC-1 and PC-2) derived from backbone torsion angles. (b) Distance map, and (c) contact map showing the average pairwise *Cα* − *Cα* distances and contact probability across all frames. (d) Distribution of the radius of gyration (*R_g_*) for the full dataset, training subset, and DDPM-generated data. (e-g) Secondary structure probabilities for Coil, *α*-Helix, and *β*-Sheet as functions of residue indices. The plots demonstrate that the DDPM-generated conformations closely reproduce the structural properties of the original data, capturing both global and local features with high accuracy. Slight deviations in absolute secondary structure probabilities are observed for a few residues, but overall trends remain consistent.

Similarly, we analyzed the distance map (Figure 7(b)), contact map (Figure 7(c)), radius of gyration distribution (Figure 7(d)), and secondary structure probabilities (Figures 7(e-g)) for the IDR Ash1. These analyses reveal that the DDPM-generated conformational ensemble aligns closely with the structural properties of the original 100% MD simulation data. The distance and contact maps demonstrate good agreement, capturing both local and global structural features. While the *R_g_* distribution qualitatively captures the compactness and heterogeneity observed in the original datasets, quantitative deviation from the 100 % MD simulation data is present. While the secondary structure probabilities for Coil, *α*-Helix, and *β*-Sheet formations show trends similar to the original data, some residues exhibit deviations in absolute probabilities, particularly for *α*-Helix and *β*-Sheet. In hindsight, for intrinsically disordered proteins (IDPs) such as Ash1, deviations in secondary structure probabilities between the DDPM-generated ensemble and the full MD data, particularly in *α*-helix and *β*-sheet content–are expected and were explicitly discussed in our analysis. These discrepancies arise primarily due to the inherent conformational heterogeneity and lack of stable secondary structure in IDRs, which pose challenges for learning fine-grained structural propensities from limited data. Moreover, the DDPM tends to reproduce distributions present in the training set, which may underrepresent rare secondary structure transitions. This highlights the need for more diverse or enhanced sampling-based training datasets for disordered systems, a direction we emphasize in our concluding remarks.

#### Validation Using IDP ***α***-Synuclein

To further test the capability of DDPM in generating conformational ensembles, we investigated whether the model could accurately replicate the structural properties of a larger IDP. Specifically, we applied DDPM to *α*-Synuclein, a 140-residue IDP known for its highly dynamic and heterogeneous ensemble of conformations. Figure S10 presents the 2D FES plots along the first two principal components (PC-1 and PC-2) of the backbone torsion angle for the full MD simulation data, the training subset, and the DDPM-generated data. Remarkably, for this larger and more complex IDP, the DDPM-generated conformations effectively recapitulate the essential features of the original FES while also sampling regions sparsely populated in the training data, showcasing the model’s ability to capture the full conformational landscape. Furthermore, the generated FES landscape is also consistent with our previous analyses of FES plots obtained by training the DDPM directly on torsion angle data.

Additional analyses were performed to validate the structural properties of *α*-Synuclein. These include the distance map (Figure 8(a)), contact map (Figure 8(b)), radius of gyration (*R_g_*) distribution (Figure 8(c)), and secondary structure probabilities (Figures 8(d-f)) for Coil, *α*-Helix, and *β*-Sheet formations. The results indicate certain deviations between the DDPM-generated conformations and the fully MD-simulated dataset, particularly in the distance map, which appears to be influenced by characteristics inherent in the training data. The *R_g_* profile is generally well reproduced, except at very large values corresponding to a low-probability regime. While slight deviations in absolute probabilities were observed for *α*-Helix formation in certain residues, the overall trends are consistent with the original data. These analyses confirm that the DDPM-generated conformations accurately represent the structural heterogeneity of *α*-Synuclein, including its local and global structural properties.

**Figure 8:**
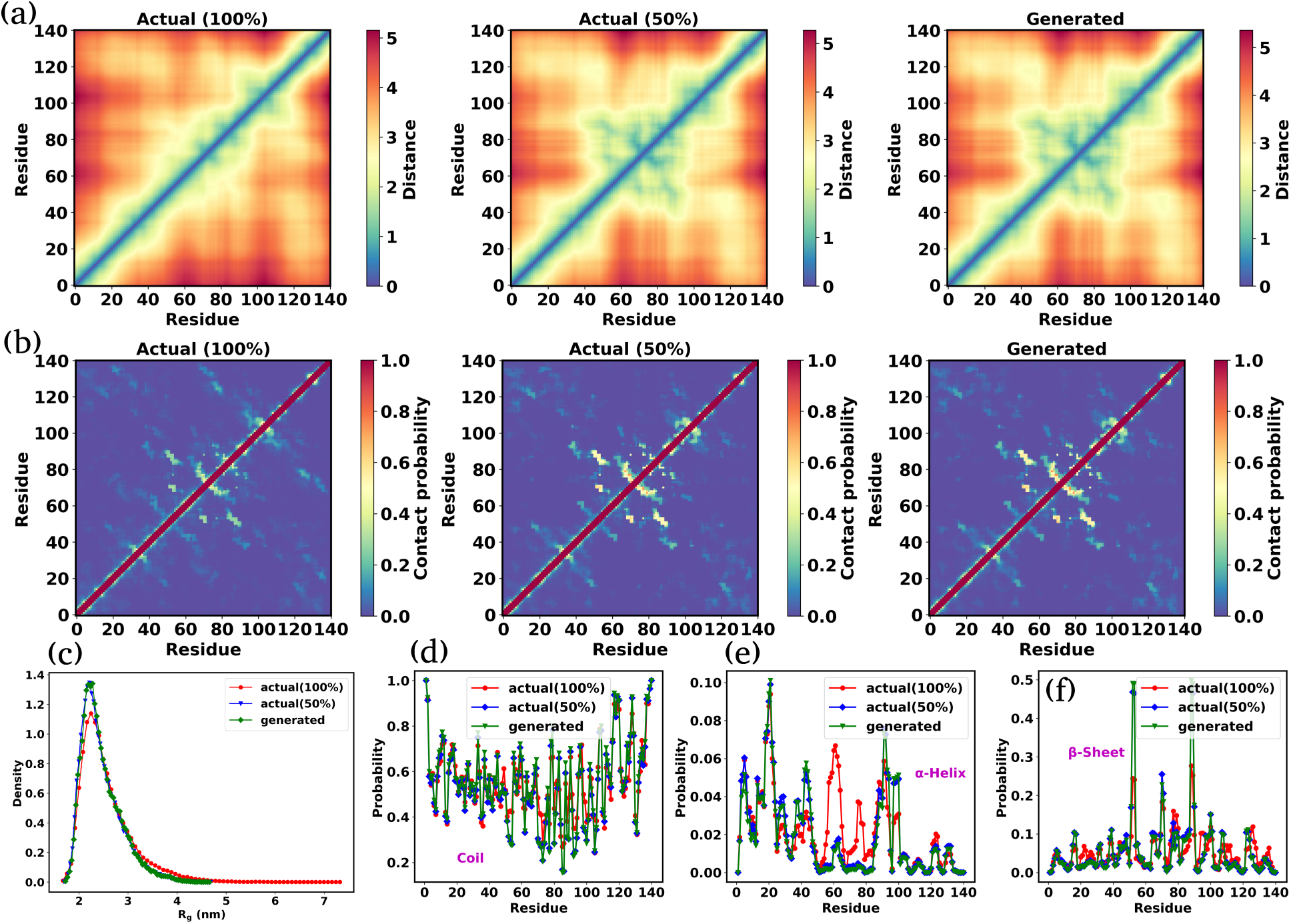
Comparison of structural features for *α*-Synuclein obtained from MD simulation, training data, and DDPM-generated ensembles. (a) Distance map and (b) contact map comparing the DDPM-generated conformations with the original MD data, showing excellent agreement in capturing local and global structural features. (c) The (*R_g_*) distribution demonstrates that the DDPM-generated conformations closely align with the compactness and heterogeneity of the original dataset. (d-f) Secondary structure probabilities for Coil, *α*-Helix, and *β*-Sheet formations, highlighting trends consistent with the original data, with slight deviations observed in *α*-Helix probabilities for certain residues. These results confirm that DDPM accurately captures the conformational heterogeneity and structural properties of *α*-Synuclein..

### Computational Performance Benchmark: MD Simulations vs. DDPM Ensemble Generation

Since the original datasets used in our work were generated by D. E. Shaw Research using the a99SB-disp force field on the specialized Anton supercomputer, we could not reproduce these simulations in full. Instead, we ran shorter benchmark MD simulations on our local HPC cluster for both a small globular protein (Trp-cage) and a large intrinsically disordered protein (IDP), *α*-Synuclein, using the same simulation setup as reported in the original studies.^79^ From these benchmarks, we estimated the MD simulation performance *P*_md_ (in ns/day) and extrapolated the total time it would take to simulate the full trajectories of 100 *µ*s for Trp-cage and 73 *µ*s for *α*-Synuclein. In parallel, we recorded the total wall-clock time required for training the DDPM model and generating an equivalent number of samples as present in the full-length MD datasets.

The comparison is summarized in Table 1. For Trp-cage, generating an equivalent dataset using DDPM required significantly less time than performing a 100 *µ*s MD simulation on our hardware. For *α*-Synuclein, the advantage is even more pronounced. Performing such a long-timescale simulation using conventional MD would have taken several years on our local cluster, making it practically infeasible. In contrast, the DDPM model was able to generate the same number of configurations within a matter of days. These observations clearly indicate that DDPM offers significant speed-ups, especially for larger, more complex systems. All molecular dynamics simulations and training of the DDPM models were conducted using an Intel(R) Xeon(R) Platinum 8352Y CPU at 2.20 GHz, along with an NVIDIA RTX A6000 GPU.

**Table 1:** Comparison of performance between MD simulations and DDPM.

| System | $P_{\text{md}}$ (ns/day) | Total MD time ($\mu s$) | Time required for MD (day) | Time required for ML (day) |
| --- | --- | --- | --- | --- |
| Trp-cage | $\sim 832.0$ | 100.0 | $\sim 120.19$ | $\sim 3.32$ |
| α-Synuclein | $\sim 35.0$ | 73.0 | $\sim 2085.71$ | $\sim 12.02$ |

### Structural Comparison of DDPM-Generated Configurations with Original Data

To visualize how closely the structure of the generated proteins aligns with the original data, we individually plotted the configurations. First, we extracted the configurations of each protein from their respective minima on the FES plots of the original dataset and used them as reference structures for calculating root-mean-square deviations (RMSDs). We computed the RMSDs of the DDPM-generated ensemble relative to these reference structures and selected five configurations for each protein with the smallest RMSD values. Subsequently, we overlaid these five DDPM-generated configurations with their respective reference structures from the original simulation data. In all overlay figures, the reference structures are shown in blue, while the generated structures are in red. Table S1 provides the corresponding mean RMSD values, highlighting how closely the generated structures resemble the original configurations.

Figures 9(a) and (b) show the superimposed configurations of the Trp-cage mini-protein, revealing its folded *α*-helix and unfolded random coils, respectively. Similarly, Figures 9(c-e) visualize the superimposed configurations of the folded BPTI protein. For both Trp-cage and BPTI, the DDPM-generated configurations closely match the secondary structure observed in the reference data. Figures 9(f-g) and (h-j) present the overlaid structures of the IDR Ash1 and IDP *α*-Synuclein, respectively. These visualizations confirm that the DDPM-generated configurations not only closely replicate the original IDP structures but also effectively capture their significant conformational heterogeneity.

**Figure 9:**
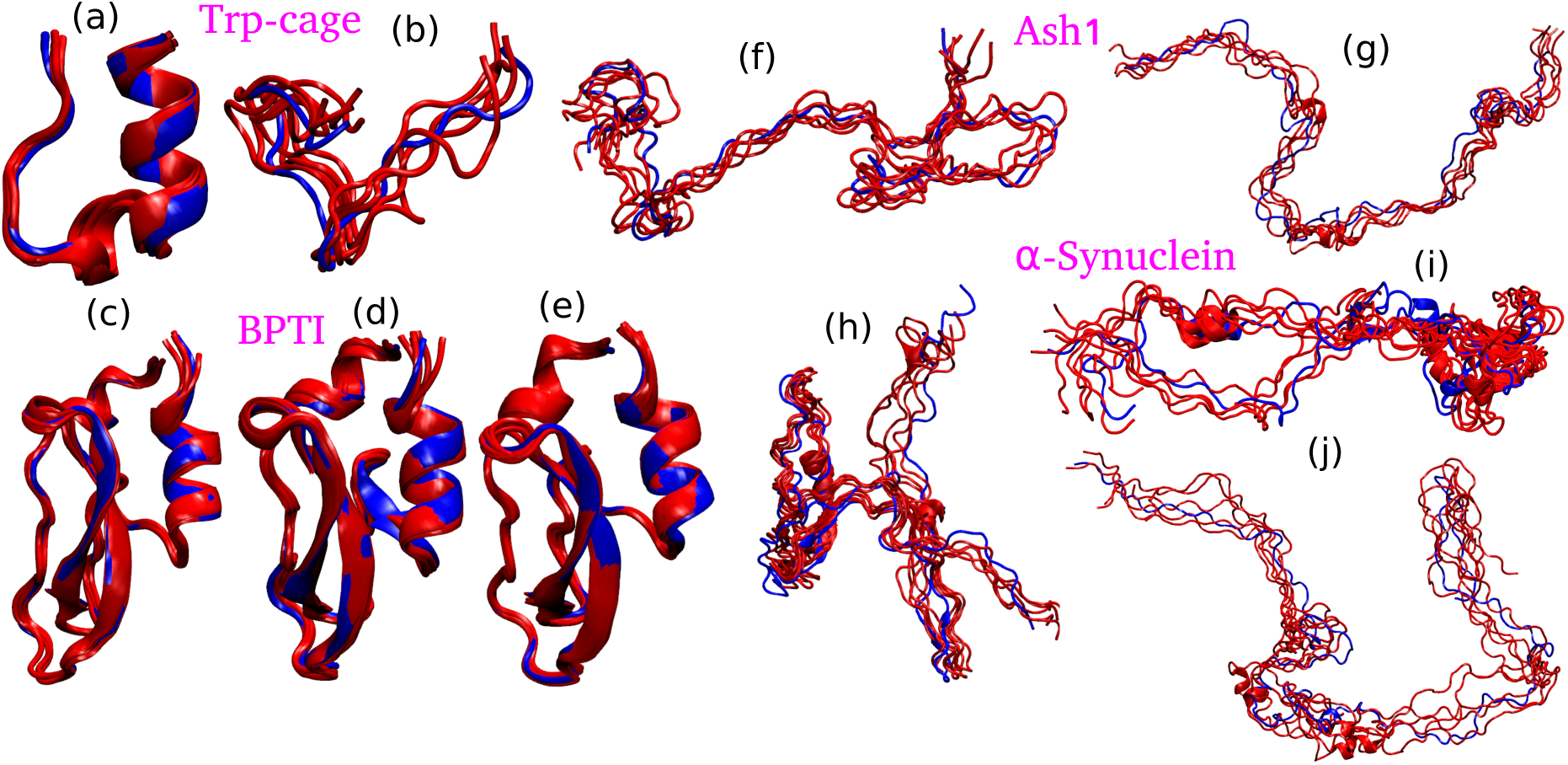
Overlay of DDPM-generated configurations (red) with reference structures (blue) extracted from the minima of FES plots of the original dataset. (a-b) Superimposed configurations of the Trp-cage mini-protein, illustrating its folded *α*-helix and unfolded random coil states, respectively. (c-e) Overlay of the folded BPTI protein configurations, showcasing the accurate reproduction of secondary structures by the DDPM-generated ensembles. (f-g) and (h-j) Superimposed configurations of IDR Ash1 and IDP *α*-Synuclein, respectively, demonstrating the ability of DDPM to closely capture the conformational heterogeneity of IDPs.

## DISCUSSION AND CONCLUSIONS

The findings presented in our study demonstrate the efficacy of the DDPM in enhancing sampling accuracy within MD simulations, ranging from small folded proteins to large IDPs. By using the principles of diffusion processes, DDPM not only generates realistic molecular conformations but also effectively samples regions of the conformational space that are sparsely populated in the training data. This capability is particularly significant when considering the challenges associated with exploring high-dimensional biomolecular landscapes.

Our study began with training DDPM on torsion angle for Trpcage mini-protein and IDP *α*-Synuclein. We noted that the FES generated by DDPM closely resembled those obtained from full-length MD simulations, even when trained on smaller subsets of data. The analysis of torsion angle distributions revealed that DDPM accurately captured complex features, including bimodal distributions. Our findings highlight two primary ways through which DDPM enhances sampling: (i) interpolation within existing data distributions and (ii) the generation of new, low-probability samples.

Expanding upon this, we trained DDPM using all-atom coordinates and observed its remarkable ability to comprehensively explore the conformational spaces of both folded proteins and IDPs. Through comparisons across a range of structural properties including free energy surfaces, inter-residue distances, contact maps, radius of gyration distributions, and secondary structure probabilities, DDPM-generated conformational ensembles demonstrated decent alignment with the original MD simulation data. These results affirm the model’s ability to generate conformations that are not only statistically representative but also structurally relevant.

Moreover, we deliberately minimized hyperparameter tuning by employing a common model architecture across all systems, with only minor adjustments based on input representation (e.g., torsional versus Cartesian features), as described in the Supplementary Information. This restrained optimization strategy ensured that the overall computational overhead remained low, preserving the practical benefits of DDPM-based sampling in terms of time and resource efficiency.

Our investigation highlights both the strengths and limitations of DDPM-generated ensembles in biomolecular conformational sampling. While DDPM effectively captures key structural features and enhances sampling in sparsely populated regions of the free energy landscape, it also exhibits certain biases and limitations. Specifically, the model can fail to sample certain low-probability regions, and its performance is strongly influenced by the quality and representativeness of the training data segments. This effect is particularly evident in more flexible proteins such as Trp-cage and intrinsically disordered proteins (IDPs), where conformational variability is high, compared to BPTI, which is structurally more rigid and displays greater agreement between DDPM-generated and MD-simulated conformations. Notably, in the case of IDPs, deviations in secondary structure propensities between the generated and reference datasets suggest that DDPM may not fully capture the complex conformational heterogeneity of disordered proteins.

Nevertheless, DDPM offers significant advantages in certain aspects of conformational sampling. The model is capable of generating states in low-probability regions that were either sparsely sampled or entirely absent in conventional MD trajectories. Furthermore, it effectively interpolates transitions between well-separated basins, providing insights into conformational changes that may be difficult to observe within finite-length MD simulations. However, while these newly generated states suggest the potential for uncovering novel biomolecular conformations, independent validation is required to establish their physical relevance.

In hindsight it is also important to recognize that even “100% simulated data” has inherent limitations. Since MD simulations are finite in length and constrained by initial conditions, their ability to comprehensively sample the conformational landscape is not absolute. This means that some of the “ground-truth” reference datasets used for benchmarking may themselves suffer from incomplete sampling, making it difficult to definitively assess whether deviations observed in DDPM-generated ensembles stem from model artifacts or highlight previously unexplored regions of the free energy landscape. This underscores the need for a cautious interpretation of both MD and DDPM-generated data, with an emphasis on rigorous validation using orthogonal computational or experimental approaches.

One of the key limitations of the current DDPM-based approach lies in the temporally discontinuous nature of the generated conformational ensembles. Since DDPMs generate statistically plausible conformations in an uncorrelated manner, the resultant trajectories lack the continuous time evolution characteristic of molecular dynamics (MD) simulations. This inherently prevents the extraction of kinetic information, such as transition rates or time-resolved state populations, which are essential for understanding the dynamic behavior of biomolecular systems. While the generated ensembles may approximate equilibrium distributions, they do not preserve dynamical correlations, thus precluding their direct use in constructing kinetic models like Markov state models (MSMs) or in estimating experimentally measurable timescales.

Additionally, the DDPM used in this study is currently non-transferable, requiring a dedicated model to be trained for each protein of interest. While this approach allows for system-specific optimization and resolution, it limits broader applicability and scalability. Recent developments in generative modeling of IDPs have introduced transferable frameworks that can generate conformational ensembles from primary amino acid sequences alone.^72,73^ However, these models are predominantly based on coarse-grained representations or trained on simulations employing implicit solvent models, and as such, do not offer atomistic detail or capture solvent-mediated interactions that are often critical for IDP function. Achieving transferability in the context of atomistic, explicit-solvent models would represent a significant step forward but would require a substantial amount of equilibrated simulation data across a diverse set of protein sequences. Such a task is computationally intensive, particularly for large and heterogeneous IDPs, and would necessitate considerable investment in dataset curation, model architecture design, and high-performance computing resources. Nevertheless, the development of a transferable atomistic DDPM remains a promising direction, potentially enabling general-purpose generative tools for biophysical modelling.

Our present study is also constrained by the thermodynamic conditions under which the training data were generated. Specifically, the MD simulations used to train the DDPM were performed under fixed physical conditions, typically constant temperature, pressure, and solvent environment, which limits the generalizability of the generated ensembles to those specific settings. As such, the current DDPM can only sample conformations reflective of the thermodynamic ensemble present in the training dataset. A natural extension of this work would be to develop models capable of condition-dependent ensemble generation, by training on datasets spanning a range of physiological or experimental conditions.^74,81^ This would enable the generation of context-sensitive conformational ensembles and facilitate the study of condition-induced structural transitions, such as thermal unfolding or allosteric regulation.

Beyond protein conformational dynamics, we envision a broader scope for DDPMs in modeling increasingly complex biomolecular systems. One important direction is the application to protein-ligand interactions, where generative models could be used to sample binding-competent conformers or aid in pose prediction.^82–85^ Another compelling domain is the study of membrane-associated IDPs, where conformational heterogeneity is further complicated by lipid interactions and curvature sensitivity.^86,87^ Lastly, the modeling of IDP aggregation and phase behavior presents an exciting frontier, where DDPMs could be employed to explore early aggregation intermediates or sequence-dependent phase separation phenomena.^88–90^ These extensions would not only demonstrate the versatility of diffusion models in biomolecular contexts but could also provide mechanistic insights into problems of biomedical relevance.

While our results demonstrate that DDPMs can capture major features of the conformational landscape and occasionally generate novel conformers, they also highlight specific shortcomings in sampling low-probability regions that are poorly represented in the training data. This limitation poses a dual challenge: first, it constrains the models ability to resolve the fine-grained topology of the free energy surface, and second, it introduces conformational states whose physical validity is uncertain in the absence of independent verification. To address these issues, our future work will explore two complementary strategies. First, we aim to incorporate data from enhanced sampling methods such as metadynamics or replica-exchange MD to increase representation of rare events, while exercising caution to avoid biasing the learned distributions. Second, we plan to develop conditional variants of DDPMs, wherein generation can be guided by predefined collective variables or thermodynamic constraints, enabling focused sampling of sparsely populated yet functionally important regions of conformational space. Together, these directions promise to further enhance the utility of DDPMs in advancing our understanding of protein dynamics and function.

In summary, this study offers a critical evaluation of DDPMs for generative modeling of protein conformational dynamics across both structured and intrinsically disordered systems. While the models successfully reproduce dominant thermodynamic features from limited MD data, they underperform in capturing low-populated conformers, an inherent limitation rooted in training data sparsity. Rather than presenting only the strengths, we have deliberately highlighted these shortcomings to provide an objective baseline for the field. In the fast moving field of machine learning in molecular sciences, we believe such candid assessment is essential for guiding future innovations. Our findings underscore the need for more transferable architectures, integration with enhanced sampling data, and uncertainty-aware modeling to better explore the subtle yet critical fringes of biomolecular conformational space.

## ASSOCIATED CONTENT

### Data and code availability

All data supporting the findings are included within the manuscript. Additionally, the code and comprehensive documentation for training the DDPM can be publicly accessed on GitHub via the following link: https://github.com/palash892/DDPM_conformational_ ensemble

### Supporting Information

The Supporting Information (SI) provides additional figures (S1-S9), containing Free energy surface (FES) plots of the *α*-Synuclein along the first two principal component, Ramachandran plots for Trp-cage, *α*-Synuclein, Probability distributions for selected dihedral angles in Trp-cage mini-protein,Probability distributions for selected dihedral angles in *α*-Synuclein.,FES plots for the Trp-cage mini-protein along the first two principal components, Kullback-Leibler divergence between generated and actual torsion angle distributions as a function of training epoch, table (S1) containing RMSD of different proteins, and supplementary methods with technical details (reparameterization scheme in DDPM, U-Net Architectures), that support the main findings presented in the manuscript.

## Acknowledgments

We acknowledge support of the Department of Atomic Energy, Government of India, under Project Identification No. RTI 4007. We sincerely acknowledge Tata Institute of Fundamental Research Hyderabad, India for providing the support of computing resources. We thank to D. E. Shaw Research for providing us the long MD simulation trajectories of Trp-cage, BPTI, Ash1, and monomeric *α*-Synuclein.^10,79^ JM acknowledges Core Research grants provided by the Department of Science and Technology (DST) of India (CRG/2023/001426).

