## Supplementary figures for "How good is Generative Diffusion Model for Enhanced Sampling of Protein Conformations Across Scales and in All-atom Resolution?"

#### Supplementary methods

##### Brief overview of reparameterization scheme in DDPM

The forward diffusion process is described by the conditional probability:

$$Q\left(x_t \mid x_{t-1}\right) = \mathcal{N}\left(x_t; \sqrt{1-\beta_t}x_{t-1}, \beta_t \mathbf{I}\right) \tag{1}$$

where the noisy data at timestep $t$ is sampled from a Gaussian distribution with mean $\mu_t = \sqrt{1-\beta_t}x_{t-1}$ and varience $\sigma_t^2 = \beta_t$. This can be expressed as:

$$x_t = \sqrt{1-\beta_t}x_{t-1} + \sqrt{\beta_t}\epsilon_{t-1} \tag{2}$$

where $\epsilon \sim \mathcal{N}(0, \mathbf{I})$ and $x_{t-1}$ is the sample at previous timestep. Now this equation suggest that the sample at each timestep conditioned on the samples from previous timesteps. To obtain a noisy sample at timestep $t$, the process iteratively evolves from $t = 0$ to $t = t - 1$. By defining $\alpha_t = 1 - \beta_t$, and $\bar{\alpha}_t = \prod_{i=1}^{T} \alpha_i$, the above equation can be rewritten as:

$$\begin{aligned} x_t &= \sqrt{\alpha_t} x_{t-1} + \sqrt{1-\alpha_t}\epsilon_{t-1} \\ \Rightarrow x_t &= \sqrt{\alpha_t}\left(\sqrt{\alpha_{t-1}} x_{t-2} + \sqrt{1-\alpha_{t-1}}\epsilon_{t-2}\right) + \sqrt{1-\alpha_t}\epsilon_{t-1} \\ &= \sqrt{\alpha_t \alpha_{t-1}} x_{t-2} + \underbrace{\sqrt{\alpha_t\left(1-\alpha_{t-1}\right)}\epsilon_{t-2}}_{\text{1st}} + \underbrace{\sqrt{1-\alpha_t}\epsilon_{t-1}}_{\text{2nd}} \end{aligned}$$

Here, the 1st and 2nd terms are Gaussian random variables with mean 0 and standard deviations $\sqrt{\alpha_t\left(1-\alpha_{t-1}\right)}$ and $\sqrt{1-\alpha_t}$ respectively. Using the property of Gaussian distributions, the sum of these two random variables is also Gaussian, with mean $\mu = \mu_1 + \mu_2$ and variance $\sigma^2 = \sigma_1^2 + \sigma_2^2$. By using this property, we can write the above equation as:

$$\begin{aligned} x_t &= \sqrt{\alpha_t \alpha_{t-1}} x_{t-2} + \sqrt{\alpha_t\left(1-\alpha_{t-1}\right) + \left(1-\alpha_t\right)} z_{t-2} \\ &= \sqrt{\alpha_t \alpha_{t-1}} x_{t-2} + \sqrt{1-\alpha_t \alpha_{t-1}} z_{t-2} \\ \Rightarrow x_t &= \sqrt{\bar{\alpha}_t} x_0 + \sqrt{1-\bar{\alpha}_t}\epsilon \end{aligned}$$

From this, the probability distribution of $x_t$ conditioned on $x_0$ becomes:

$$Q\left(x_t \mid x_0\right) = \mathcal{N}\left(x_t; \sqrt{\bar{\alpha}_t} x_0, \left(1-\bar{\alpha}_t\right)\mathbf{I}\right) \tag{3}$$

This formulation, often referred to as the reparameterization trick, allows sampling at any arbitrary timestep $t$ without the need to iterate over all prior timesteps. For further details, refer to the blog post by Nain.[1]

#### Adapting U-Net Architectures Based on Input Feature Complexity

The complexity of the input features (e.g., torsion angles or all-atom coordinates) determines the number of layers in the U-Net, enabling more accurate data generation. In our code, the parameter `dim_mults` controls the number of downsampling and upsampling blocks in the U-

Net architecture. For torsion angle data from the Trp-cage mini-protein and $\alpha$-Synuclein, we used the same architecture, with `dim_mults` set to $(1, 2, 4, 8)$. This configuration corresponds to the architecture shown in Figure 1 of the main draft. When working with all-atom coordinate data, the architecture varies based on the system. For Trp-cage, BPTI, and Ash1, we used `dim_mults` $= (1, 2, 4, 8, 16, 32)$. In the case of $\alpha$-Synuclein, we selected `dim_mults` $= (1, 2, 4, 8, 16, 32, 64)$.

#### Supplementary figures

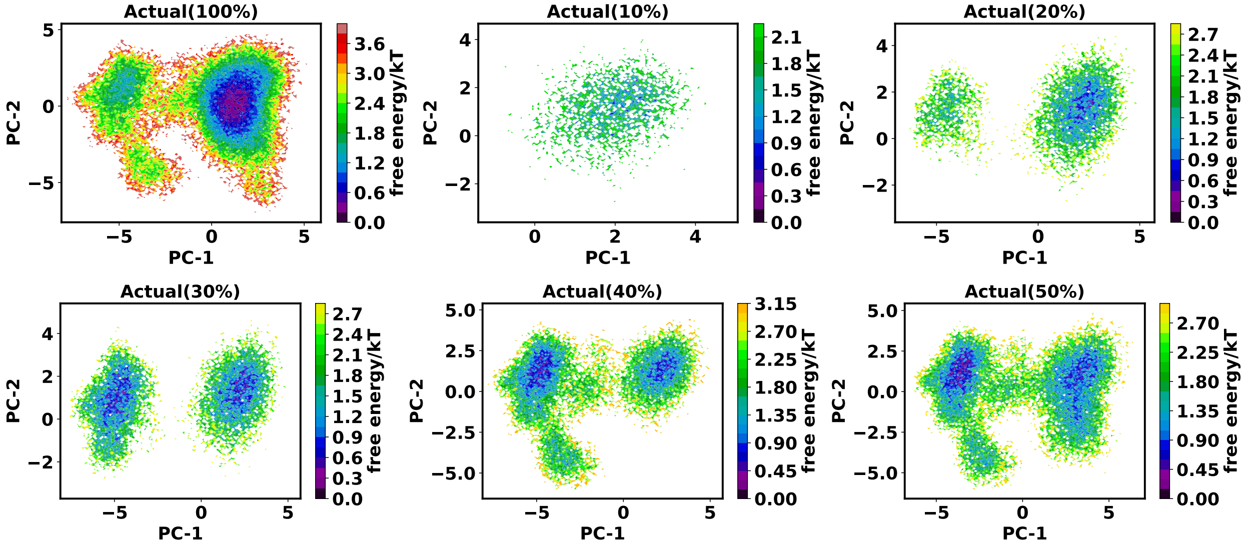


Figure S1: Free energy surface (FES) plots of the $\alpha$-Synuclein along the first two principal component, computed from various percentage of actual data.

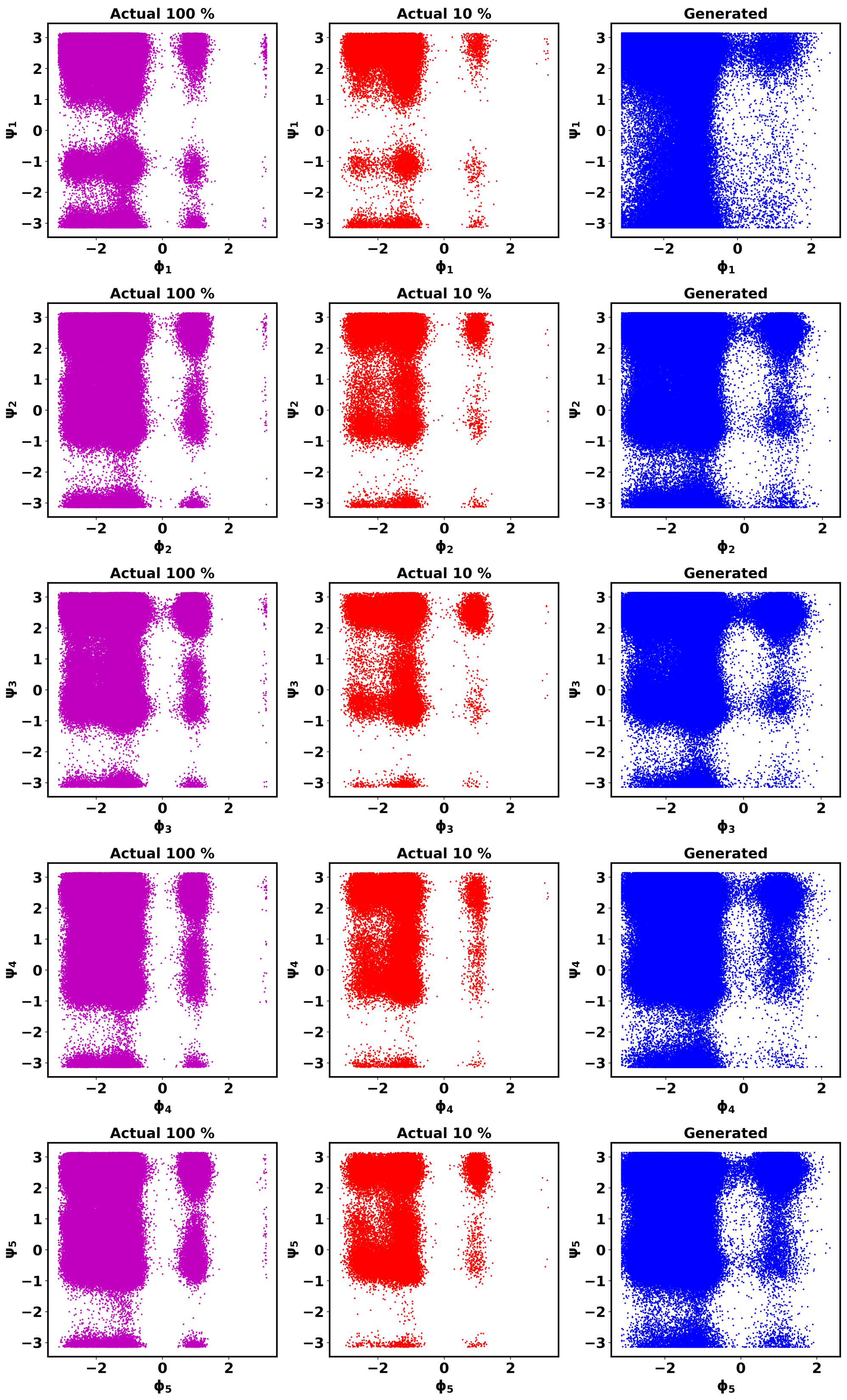


Figure S2: Ramachandran plots for the $\phi - \psi$ combinations in Trp-cage

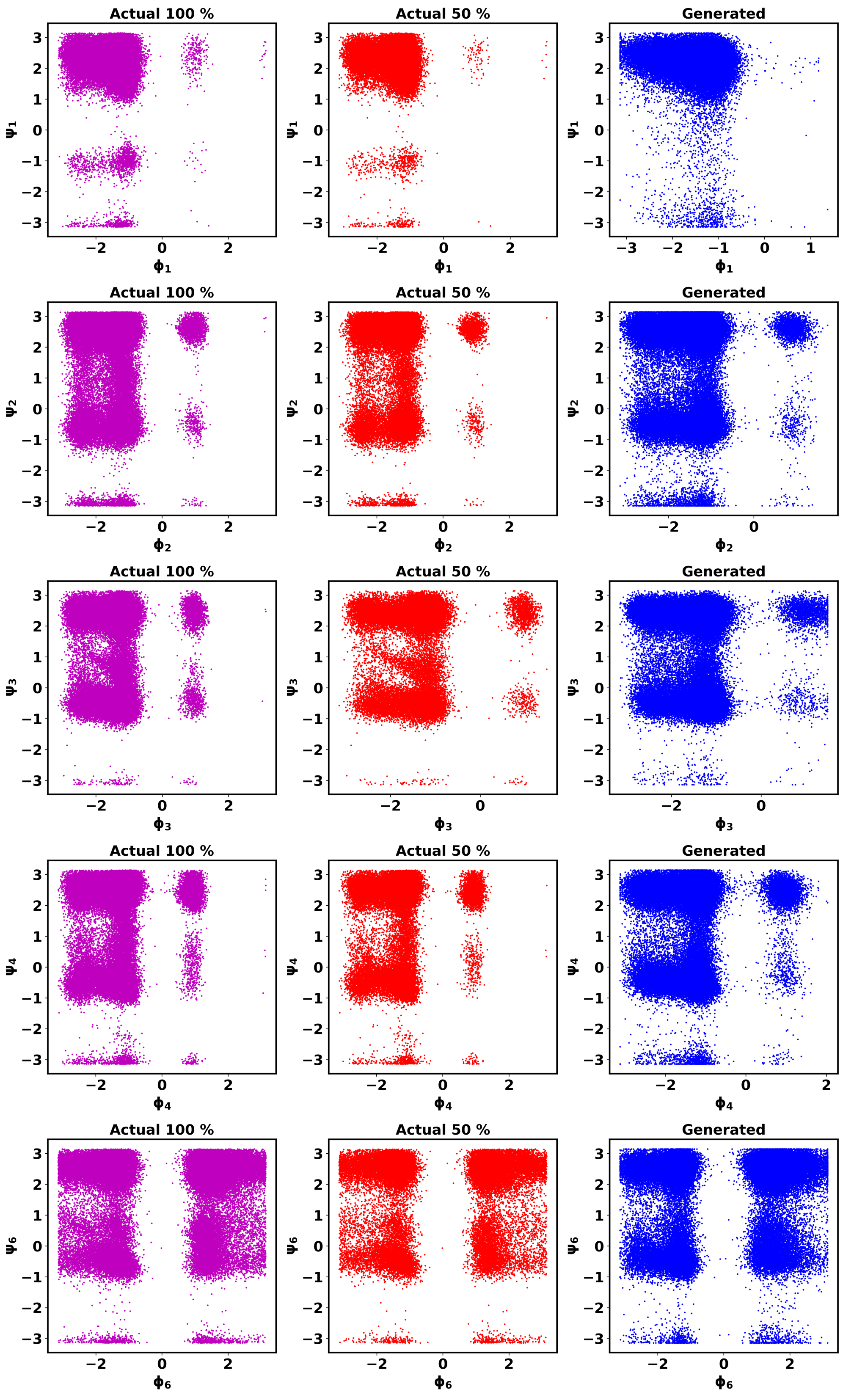


Figure S3: Ramachandran plots for the $\phi - \psi$ combinations in $\alpha$-Synuclein

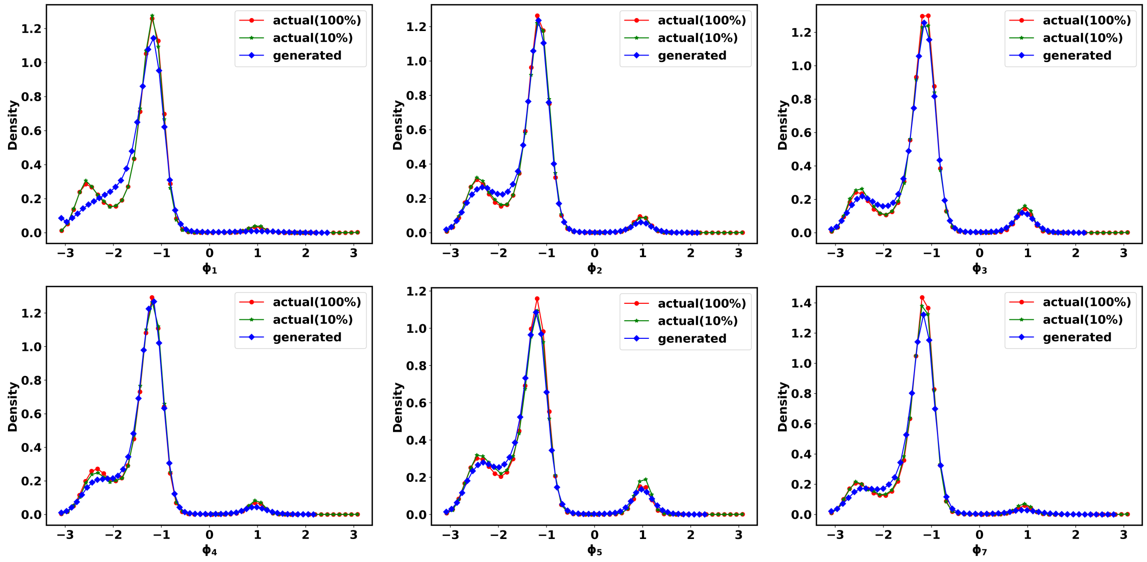


Figure S4: Probability distributions for selected dihedral angles in Trp-cage mini-protein

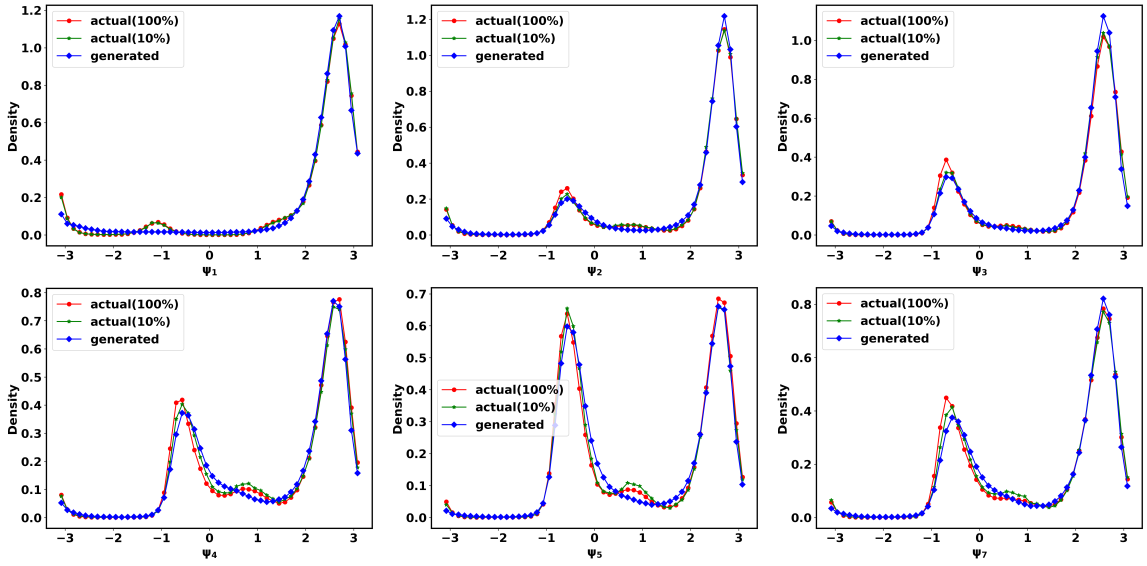


Figure S5: Probability distributions for selected dihedral angles in Trp-cage mini-protein

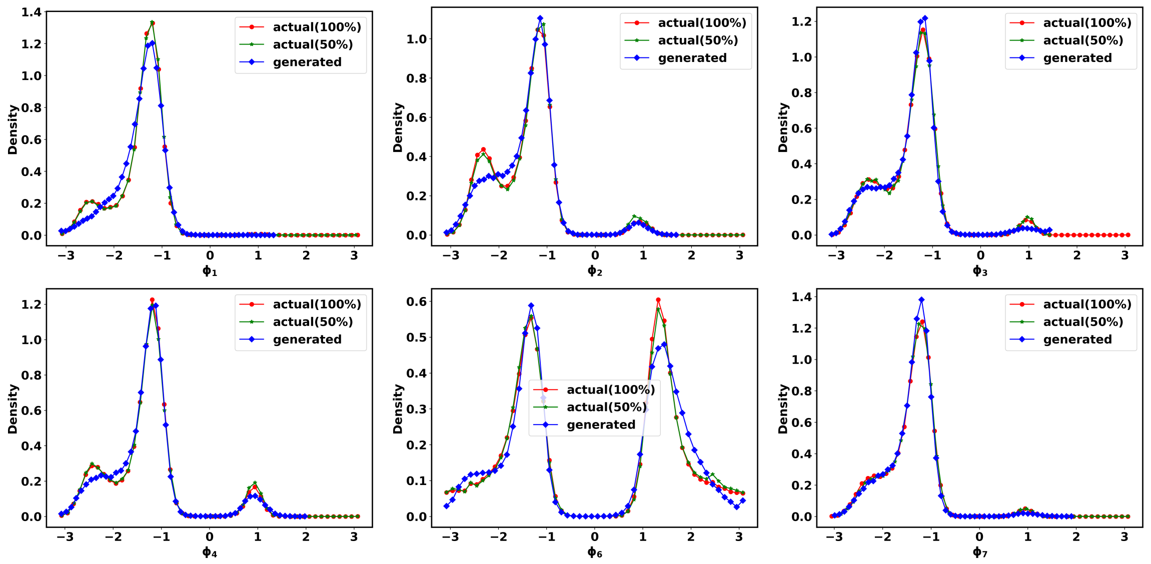


Figure S6: Probability distributions for selected dihedral angles in $\alpha$-Synuclein.

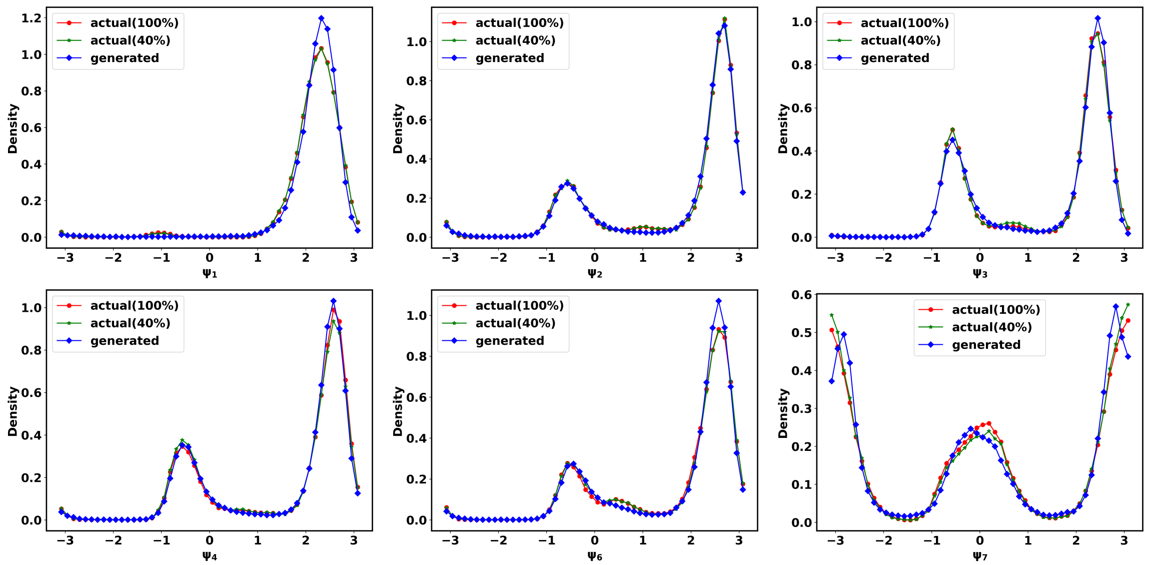


Figure S7: Probability distributions for selected dihedral angles in $\alpha$-Synuclein.

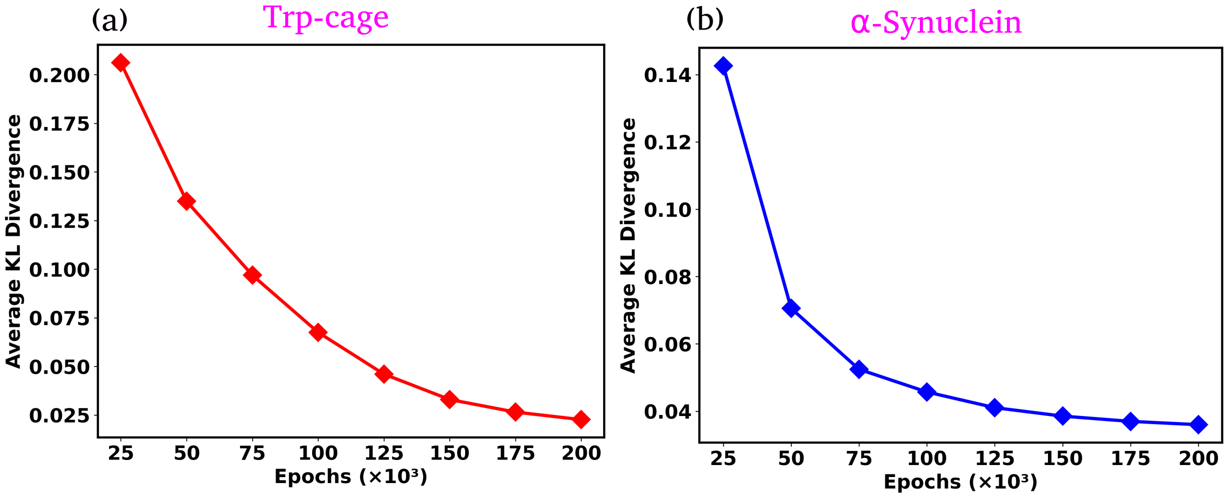


Figure S8: KL divergence between the true and generated torsion angle distributions as a function of training epochs for Trp-cage mini-protein (a) and $\alpha$-Synuclein, respectively. The plots show a clear monotonic decrease in KL divergence over time, indicating that the generated samples increasingly resemble the true distribution as training progresses.

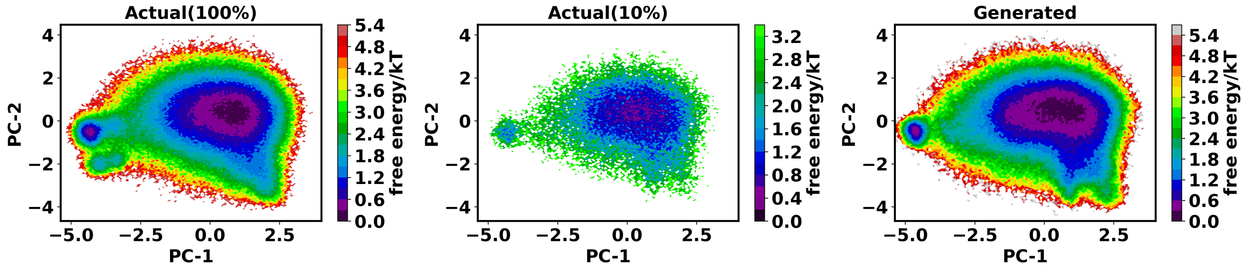


Figure S9: FES plots for the Trp-cage mini-protein along the first two principal components (PC-1 and PC-2) are shown for the full 100% MD simulation data, the training subset, and the DDPM-generated data. Here we calculated the backbone torsion angles from the all-atom coordinate itself.

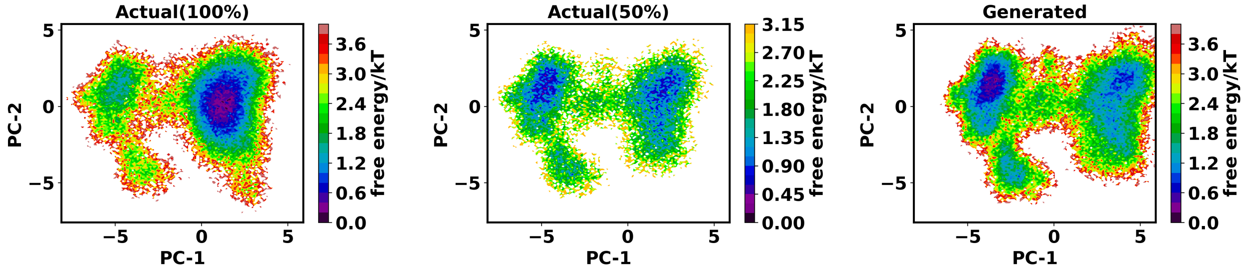


Figure S10: FES plots for the IDP $\alpha$-Synuclein along the first two principal components (PC-1 and PC-2) are shown for the full 100% MD simulation data, the training subset, and the DDPM-generated data. Here we calculated the backbone torsion angles from the all-atom coordinate itself.

Table S1: RMSD for different system conformations.

| **System** | **Conformation** | **Mean RMSD (nm)** | **STD in RMSD (nm)** |
|---|---|---|---|
| Trp-cage | a | 0.0828 | 0.0028 |
| Trp-cage | b | 0.3459 | 0.0124 |
| BPTI | c | 0.0987 | 0.0034 |
| BPTI | d | 0.1062 | 0.0015 |
| BPTI | e | 0.0893 | 0.0031 |
| Ash1 | f | 0.3396 | 0.0544 |
| Ash1 | g | 0.5185 | 0.0248 |
| $\alpha$-Synuclein | h | 0.4300 | 0.0676 |
| $\alpha$-Synuclein | i | 0.5224 | 0.05576 |
| $\alpha$-Synuclein | j | 0.6990 | 0.0888 |

### References


(1) Aakash Nain. A deep dive into ddpms. URL https://magic-with-latents.github.io/latent/posts/ddpms/part3/.
